# From hotspot dependence to distributed robustness in resistance-aware lead optimization

**DOI:** 10.64898/2026.06.16.732538

**Authors:** Yuxuan Wang, Bowen Xiao, Jingyi Kang, Huizi Cui, Yu Fu, Wen Li, Silvio E. Perea, Weiwei Han

## Abstract

Drug resistance remains a recurrent failure mode in targeted anticancer and antiviral therapy, and resistance evidence often enters only after compound selection. ResistAgent is an evidence-constrained framework that converts mutational liabilities into design-time objectives through site- and combo-aware resistance mapping, deterministic mechanism diagnosis and robust counter-design. In EGFR-Erlotinib and HIV-RT-Rilpivirine, the framework separated residue-level liabilities from observed HIV combination liabilities and linked prioritized mutations to anchor loss, pocket rearrangement, electrostatic shifts and contact redistribution. Same-budget paired searches showed that robust objectives changed lower-tail mutant-panel behavior and interaction-dependence profiles while prioritizing robustness over average-affinity behavior. Under predefined liability panels, robust-objective trajectories shifted support away from mutable hotspot contacts toward more distributed interaction networks. Supplementary physical summaries and ranking-first benchmarks support the validation envelope of this resistance-aware design strategy.

## Introduction

Resistance is a dominant attrition mode for targeted anticancer and antiviral therapies, because treatment imposes selective pressure on heterogeneous and evolvable molecular systems [1–4]. In EGFR-mutant lung cancer, kinase-domain substitutions and deletions can alter inhibitor response, and secondary changes such as T790M became canonical examples of acquired resistance under first-generation inhibitors [5–14]. In HIV therapy, reverse-transcriptase mutations can appear as single substitutions or as structured combinations, especially under non-nucleoside reverse-transcriptase inhibitor exposure [15–25].

Large public resources and structural tools have made resistance evidence increasingly accessible, including mutation databases, protein structures, ligand bioactivity repositories and docking or interaction-profiling engines [26–42].

These resources are powerful but remain fragmented: priors describe which mutations are observed, structural models describe how binding may change, and medicinal-chemistry rules describe which edits are plausible. In many practical lead-optimization settings, those signals remain annotations attached after a molecule has already been selected, so the design decision itself is still governed mainly by WT or average-affinity behavior.

The methodological gap is therefore not simply the absence of another resistance database or another docking score. The gap is the absence of a design object that combines likelihood of mutation, expected binding damage and biological tolerance early enough to influence compound birth. A mutation that is common but structurally irrelevant should not dominate the design objective, whereas a rare but highly disruptive and tolerated mutation may be worth carrying into the optimization panel. A resistance-aware design framework must make that tradeoff explicit.

The chemical consequence of this gap is hotspot dependence. A candidate may appear acceptable against the WT target while relying heavily on a small set of mutable pocket contacts; once those contacts are perturbed, the binding network can fail even if the mean panel score remains respectable. In this study, distributed robustness means that the design objective asks whether support can be less concentrated on the predefined mutable hotspot panel and more distributed across contacts that are less damaged by the selected liabilities.

The gap is particularly acute when site-level and combo-level liabilities are mixed into a single ranking. A high-risk HIV combination is not equivalent to two independent single-site liabilities, because the observed combination may encode a specific evolutionary and structural context. Conversely, a single EGFR mutation can be important because it combines appearance, structural impact and fitness tolerance. A resistance-aware design strategy therefore needs to ask three connected questions: which mutations should be considered, how they disrupt binding, and how molecular edits can reduce dependence on mutable hotspots.

This distinction changes how evidence should be represented. Site-level maps answer which mutable positions should be kept visible during optimization, whereas combo-level panels preserve observed multi-mutation objects that may carry epistatic or treatment-history information. Collapsing both regimes into a single residue heatmap would make the HIV-RT-Rilpivirine case look simpler than it is, and it would obscure why a molecule optimized only against individual sites can remain fragile against observed combinations.

ResistAgent addresses this problem as an evidence-constrained closed loop for liability-aware design. The language-model component organizes evidence, prioritizes supported liabilities, explains deterministic evidence and proposes constrained edit templates; numerical evidence comes from curated priors, structural docking, interaction fingerprints, MM/GBSA-style support and predefined benchmark models. The design target reduces lower-tail vulnerability and redistributes interaction dependence across a predefined site-plus-combo mutation panel.

The case design is also deliberate. EGFR-Erlotinib and HIV-RT-Rilpivirine are the two primary cases because they support the full chain from liability ranking to mechanism diagnosis and counter-design, whereas ABL1-Nilotinib is retained as an uncertainty-heavy boundary case. This asymmetry is a scientific choice, not a convenience: the paper tests whether observed or likely liabilities can be turned into design-time objectives under a constrained evidence regime, not whether every kinase or viral target can be solved at the same confidence level.

The broader claim is narrower than a future-resistance predictor and stronger than a one-off docking exercise. ResistAgent is intended to move resistance from post hoc annotation into design-time control, with site/combo separation, mechanism-grounded diagnosis and robust SAR providing the logic for why distributed interaction organization should matter. The manuscript therefore evaluates a closed scientific chain: liability proposal, structural failure diagnosis and counter-design under a robust objective, followed by supplementary physical support and ranking-first benchmark evidence that delimit the current validation envelope.

The design loop is summarized in Figure 1: WT structural context, mutation priors, observed combinations and fitness context are organized before liability mapping, mechanism diagnosis and counter-design, while deterministic engines provide numerical evidence and the language model remains a bounded reasoning layer.

**Figure 1.**
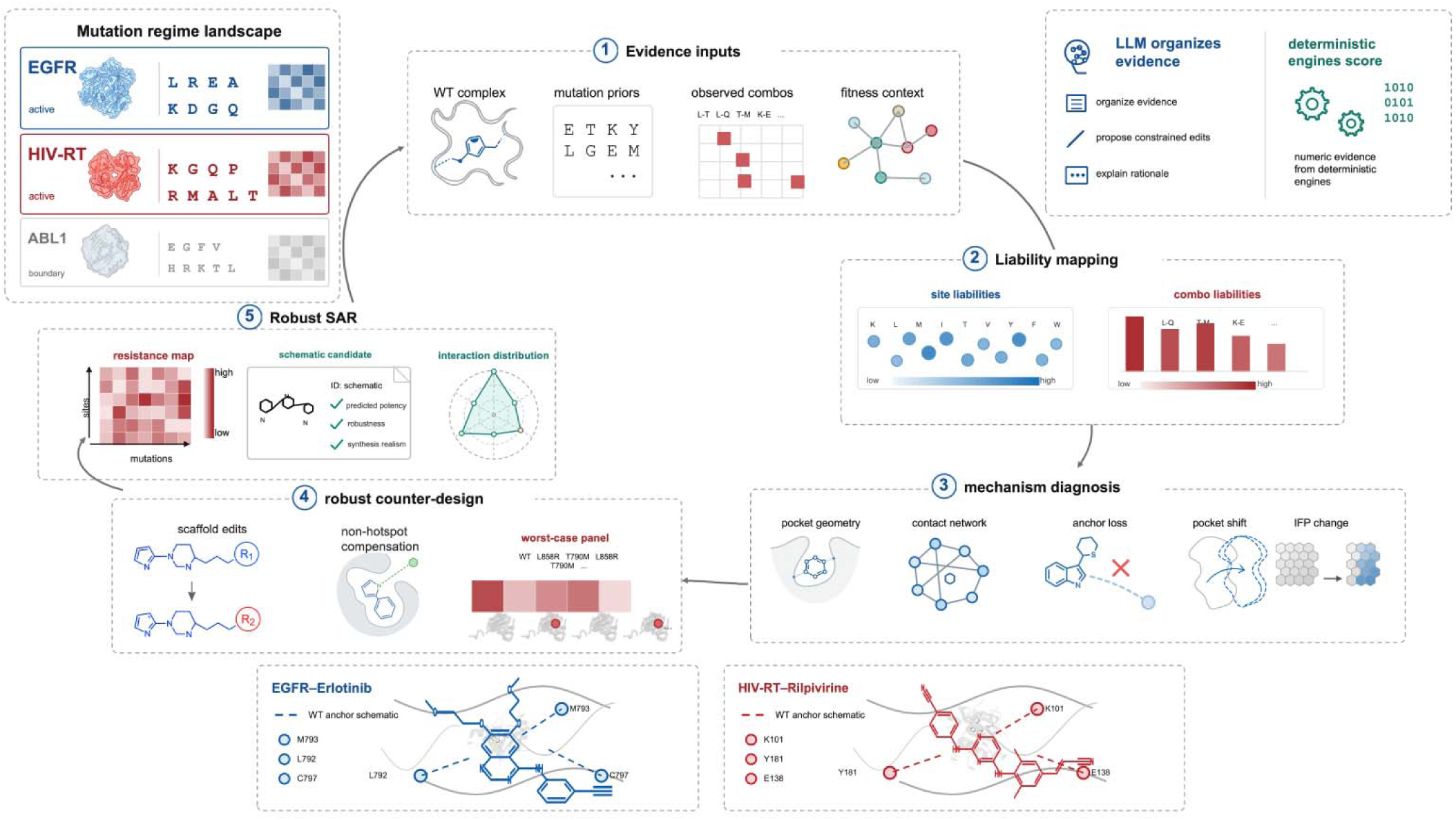
ResistAgent fronts mutational liabilities into an evidence-constrained design loop. (a) Mutation-regime landscape across EGFR, HIV-RT and boundary-case ABL1. (b) Evidence inputs combine WT complexes, mutation priors, observed combinations and fitness context. (c) The LLM organizes evidence, proposes constrained edits and explains rationale, while deterministic engines provide numerical evidence. (d) Liability mapping separates site liabilities from observed combo liabilities. (e) Mechanism diagnosis, robust counter-design and robust SAR close the loop; pocket-context examples use the same WT anchor sets carried into the structural figures.

## Results

### ResistAgent defines mutational liabilities as design-time objects

ResistAgent treats mutational liabilities as design-time objects within an evidence-constrained loop. The loop begins with WT complexes, mutation priors and observed combinations, then separates site-level and combo-level liabilities before mechanism analysis and counter-design. The primary maps retained 27 EGFR site liabilities and 3 EGFR combo liabilities, and 37 HIV site liabilities plus 16 HIV combo liabilities. This distribution establishes HIV as a site-plus-combo case (Fig. 2a,b); the ABL boundary case is reported in the Supplementary Information.

**Figure 2.**
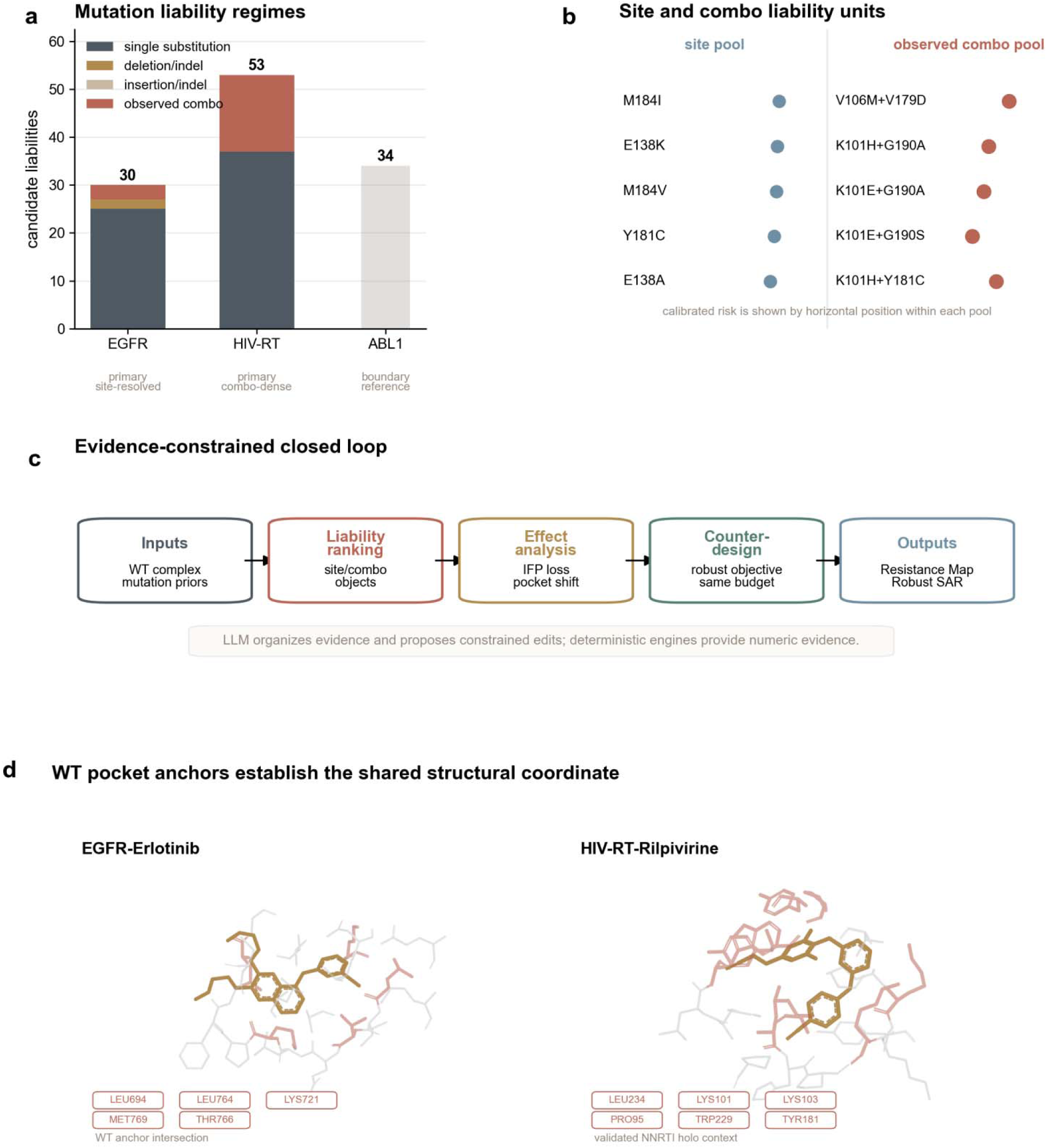
Resistance-aware design as a site- and combo-aware, evidence-constrained loop. (a) Mutation-regime landscape across EGFR-Erlotinib, HIV-RT-Rilpivirine and ABL1-Nilotinib. (b) HIV site-level liabilities are displayed separately from observed combo-level liabilities. (c) ResistAgent links deterministic inputs, mutation proposal, effect analysis and counter-design while keeping numerical evidence outside the LLM layer. (d) WT pocket anchor views for the two primary cases establish a common structural coordinate system; ABL is retained as a boundary case.

Within this boundary, the LLM layer organizes evidence and edit rationales, while deterministic engines and predefined evidence records provide docking proxies, interaction fingerprints, MM/GBSA summaries and ranking metrics (Fig. 2c). EGFR and HIV WT pocket-anchor views establish the structural coordinate used repeatedly in resistance maps, mechanism diagnosis and robust SAR (Fig. 2d).

This common coordinate matters because the same liability is seen three times: first as a prioritization problem, then as a structural failure mode, and finally as a counter-design target. The shared frame integrates scores and structures into a closed design loop.

The closed-loop framing focuses the claim on exposing mutable liabilities before design, diagnosing the interaction patterns that make them dangerous and testing whether chemistry edits can move the binding network away from the most mutable dependencies. For that reason, Resistance Map and Robust SAR are treated as the two main products of the study.

### Site- and combo-aware resistance maps identify actionable liabilities

Actionable liabilities emerged by jointly organizing appearance priors, structural impact and fitness constraints. In EGFR, the residue-level map prioritized known and structurally relevant liabilities including E746_A750delELREA and T790M together with additional candidate sites that remain within the predefined evaluation panel (Fig. 3a). Risk was treated as a calibrated product of appearance, impact and fitness terms, so the map surfaces liabilities that are actionable at design time as well as historically supported.

**Figure 3.**
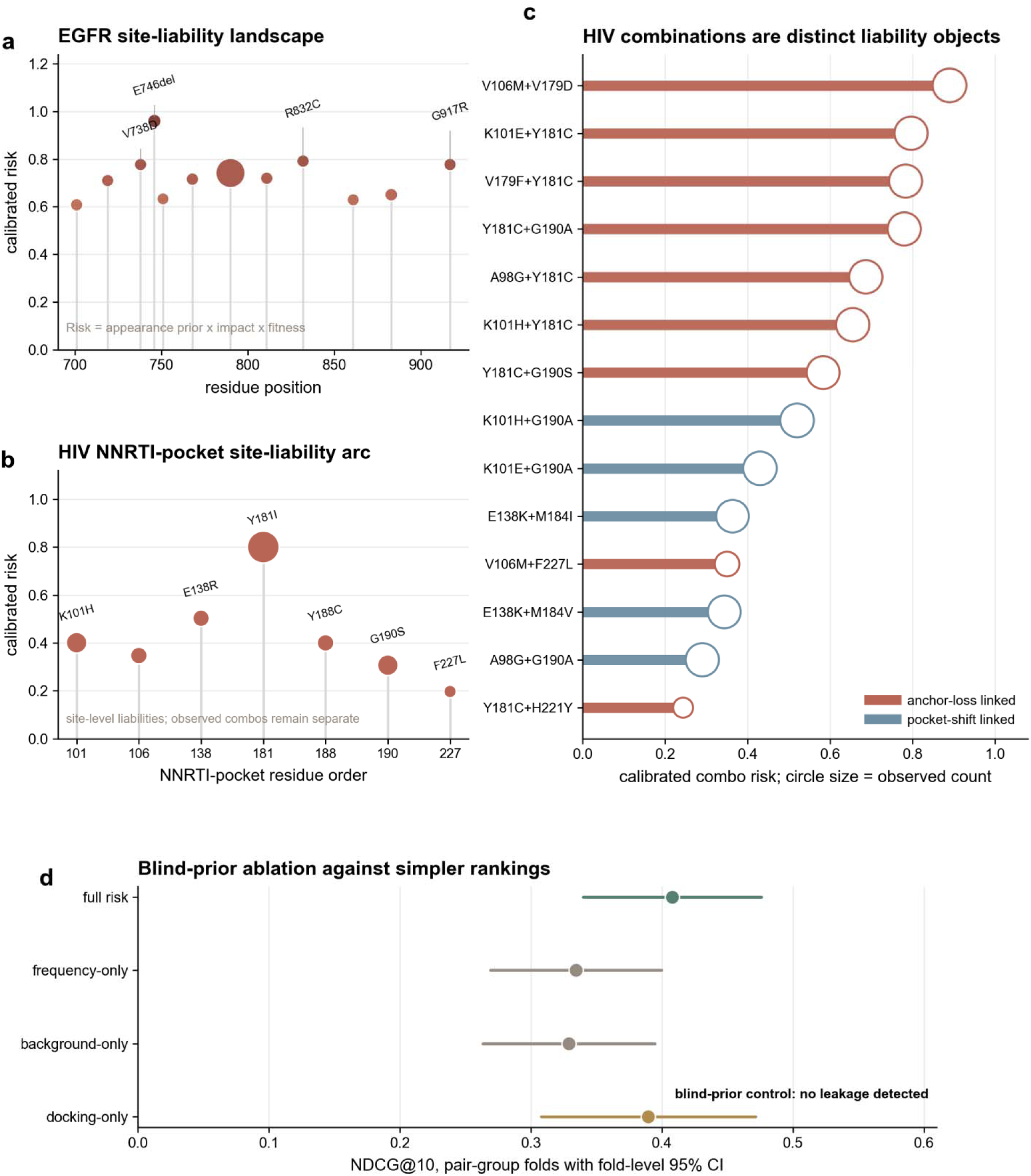
Site- and combo-aware resistance maps identify actionable mutational liabilities. (a) EGFR residue-level liability map using calibrated risk, appearance prior and structural proximity. (b) HIV NNRTI-pocket site map focused on sentinel positions. (c) Observed rilpivirine combo liabilities ranked separately from site liabilities. (d) Blind-prior ablation against simpler frequency-, background- and docking-only rankings with fold-level confidence intervals.

The EGFR map illustrates how a liability score adds design information beyond frequency. E746_A750delELREA and T790M enter the main panel through different evidence patterns: the former is represented as a high-ranked deletion liability with modeled structural support, whereas T790M carries a strong appearance term and canonical resistance relevance. Both remain useful for design because the objective keeps the most consequential mutable binding constraints visible during molecular editing.

In HIV-RT, the NNRTI pocket-centered site map highlighted sentinel residues including K101, E138, Y181, Y188, G190 and F227, while the observed combo panel ranked rilpivirine combinations such as V106M+V179D, K101E+Y181C, V179F+Y181C, Y181C+G190A and K101H+Y181C as independent combo-level liabilities (Fig. 3b,c). This separation is central to the paper: HIV combo liabilities are not synthetic afterthoughts and are not collapsed back into site-level rankings. The combo panel therefore becomes a design object in its own right, not a visual supplement to the residue map.

A blind-prior benchmark ablation further tested whether the risk organization was merely recapitulating prior frequency. The full model remained competitive against frequency-only, background-only, docking-only and structure-only alternatives under leakage-aware pair grouping (Fig. 3d). The control logic is intentionally leakage-aware: production priors can be used for case analysis, whereas blind priors are used when the question is ranking generalization. For the cancer case, sparse same-drug blind-prior evidence supports a controlled comparison against simpler rankings.

The result is a resistance map that prioritizes design-time liabilities with an explicit record of prior support. It tells the optimization procedure which mutable sites or observed combinations should be carried forward, and it establishes HIV as a site-plus-combo design problem.

### Structural effect analysis links liability ranking to failure mechanisms

The resistance maps would be insufficient if they only returned ranked mutations. ResistAgent therefore links each high-priority liability to deterministic structural evidence. WT anchor networks in EGFR and HIV-RT were defined from reference/redocking-consistent contact patterns and used as a common reference for subsequent mutant comparisons (Fig. 4a). Representative EGFR liabilities were accompanied by pocket rearrangement and interaction-pattern changes, whereas HIV examples included both a site mutant and an observed combo with anchor loss, electrostatic shift, pocket rearrangement and steric-clash signatures (Fig. 4b,c).

**Figure 4.**
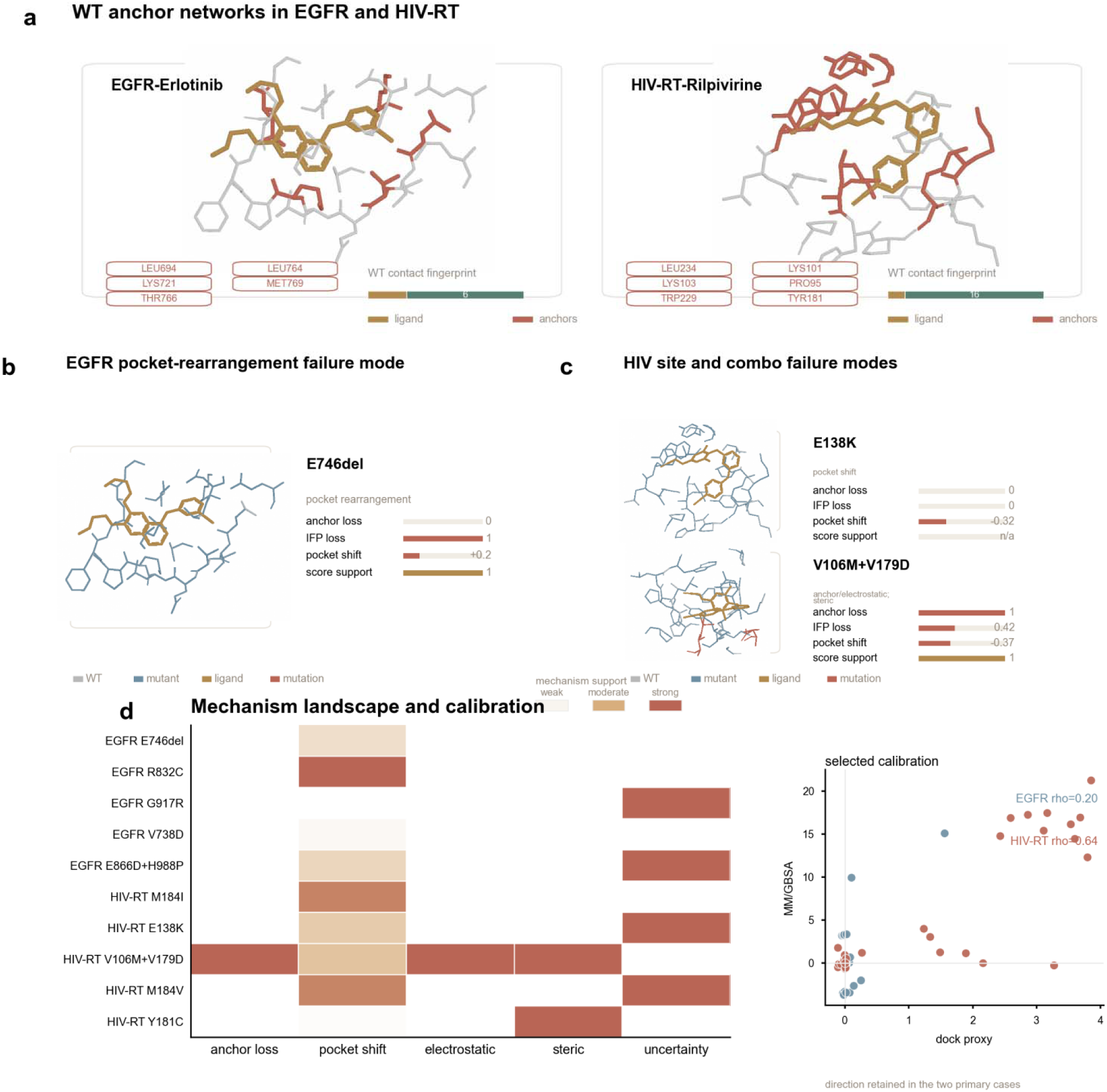
Structural effect analysis explains how top liabilities disrupt binding. (a) WT anchor networks and contact fingerprints for EGFR and HIV-RT. (b) Representative EGFR failure-mode structural summary with IFP, pocket and scoring-support readouts. (c) HIV site and observed-combo failure-mode structural summaries. (d) Mechanism landscape across top EGFR and HIV liabilities with selected calibration of docking proxy direction against MM/GBSA direction.

This step turns ranking into mechanism. The same mutation is no longer just a large score change; it is a reproducible perturbation of a specific anchor network, pocket geometry or electrostatic organization. That distinction is important because it tells the reader why a liability is risky and which part of the binding architecture is failing.

The EGFR examples show how this diagnosis works in practice. E746_A750delELREA is carried as a deletion liability with pocket-rearrangement evidence, whereas T790M+L858R combines a larger docking-proxy shift with interaction-fingerprint loss and an anchor/electrostatic signature. These examples are not treated as interchangeable score changes: one emphasizes geometric pocket remodeling, and the other illustrates how a combined kinase-domain context can disrupt the contact organization around the ligand.

The HIV examples make the site/combo distinction mechanistic. Y181C and related single-site liabilities show steric or pocket-rearrangement signatures, while top observed combinations such as V106M+V179D and K101E+Y181C add anchor-loss and electrostatic-shift patterns in the combo-level panel. Each observed combination is evaluated as a structural object with its own contact and pocket response.

Across the top liabilities, the mechanism matrix shows that pocket rearrangement is common but not exclusive; HIV combos can add anchor and electrostatic features that are not visible in a site-only summary (Fig. 4d). The calibration inset in the same panel provides support for interpreting the docking proxy as a directional effect signal in the two primary cases, while the weaker ABL calibration is reserved for the boundary-case supplement. Together, the structured readouts make failure diagnosis evidence-backed and reproducible.

### Same-budget counter-design repeats define robust objective tradeoffs

The counter-design analysis tests whether mutational liabilities can be turned into design-time objectives. The predefined robust setting combined layered interaction-fingerprint retention, mutant-panel curriculum, synthesis-realism filters, hotspot-dependence reduction and scaffold-tail diversification. Robust and naive searches used the same budget but differed in objective definition: the robust objective emphasized worst-case panel behavior, hotspot-dependence penalty and non-hotspot compensation, whereas the naive objective emphasized average-affinity behavior (Fig. 5a).

**Figure 5.**
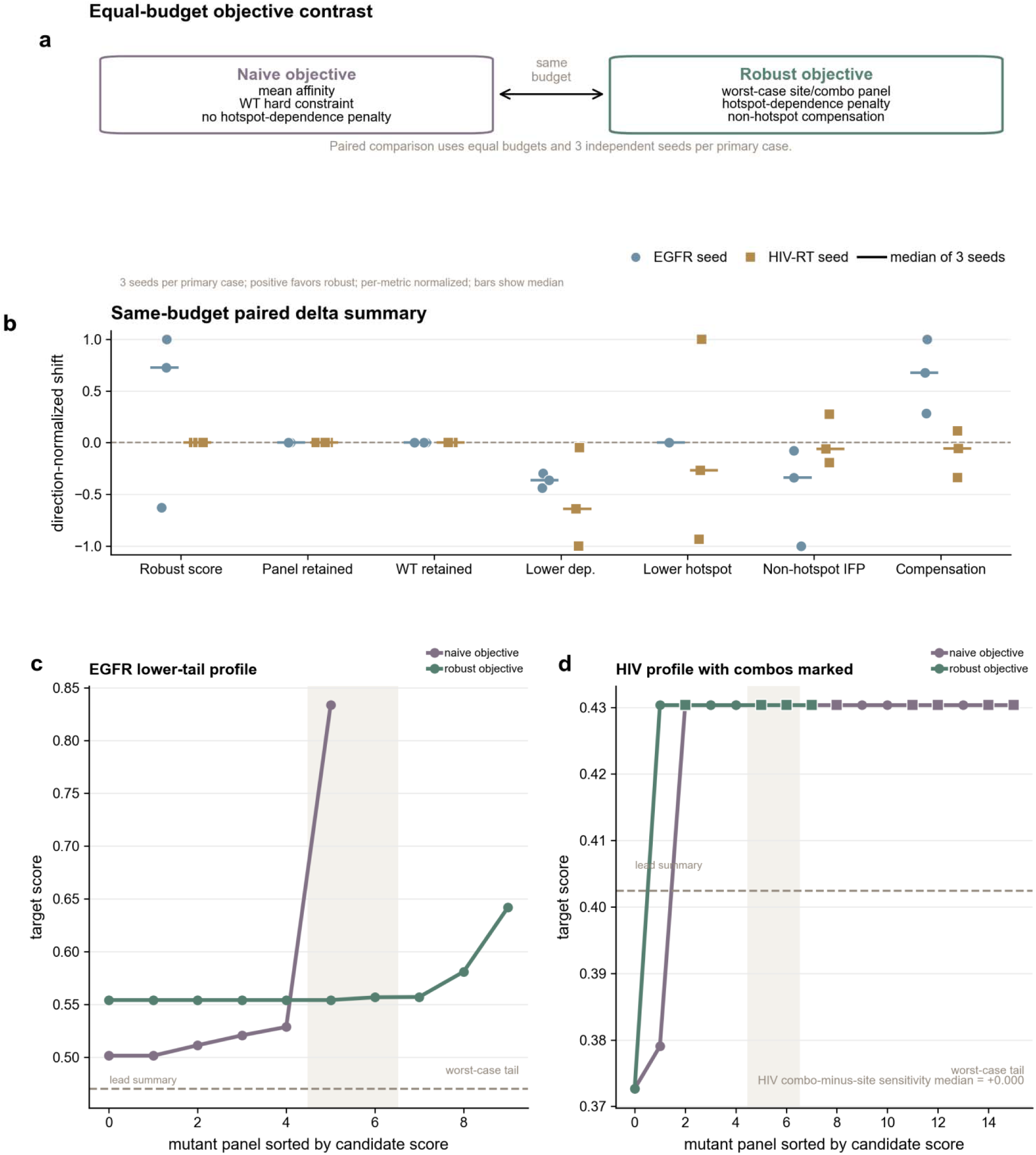
Same-budget robust-vs-naive repeats define the counter-design effect size and its limits. (a) Equal-budget robust and naive objectives evaluated across independent same-budget seeds. (b) Seed-level paired robust-minus-naive shifts across panel, target-score and dependence-rewiring metrics; positive values favor robust after applying each metric’s preferred direction. (c) EGFR mutant-panel profile for lead, naive and robust candidates. (d) HIV site-plus-combo panel profile with combo-aware evaluation.

Across three independent same-budget seeds per primary case, Figure 5 summarizes the full paired design at seed level. Robust and naive searches were matched for proposal budget, beam size, parallelization and randomization settings; the full search configuration is provided in Supplementary Table 11. In EGFR, the median robust-minus-naive shifts were +0.008 for top-20 RobustScore, +0.000 for top-20 panel passing, +0.010 for raw top-20 hotspot dependence and +0.000 for top-20 hotspot fraction. The corresponding raw top-20 dependence medians were 0.018 for robust and 0.009 for naive. In HIV-RT, the corresponding median shifts were +0.000 for top-20 RobustScore, +0.000 for top-20 panel passing, +0.017 for raw top-20 hotspot dependence and +0.009 for top-20 hotspot fraction. The raw HIV top-20 dependence medians were 0.207 for robust and 0.191 for naive. The HIV paired summary also keeps site and observed-combo sensitivity separate, with median robust-minus-naive shifts of +0.000 for the site core and +0.000 for the combo core. Because lower dependence is preferred, these repeated-seed medians do not support a uniform dependence decrease. They instead constrain the claim to objective-driven changes in lower-tail selection, compensation, diversity and dependence organization.

The paired design separates objective behavior from stochastic search variation. Robust-minus-naive shifts are therefore interpreted as properties of the objective and its evidence constraints under the matched search budget.

The repeated-seed results are interpreted as objective tradeoffs. EGFR shows a positive median shift in top-20 RobustScore and stronger objective reward for robust search, while some dependence summaries do not uniformly move in the desired direction across seeds. HIV shows site and combo cores that remain explicitly separated, with robust search changing best robust score and dependence organization under a combo-aware panel while panel-passing gains remain bounded. The main-text claim is objective-driven movement toward lower-tail and dependence-aware selection.

### Robust SAR connects objective behavior to chemistry edits

The chemical interpretation of the robust objective is summarized by the robust-SAR figure and Table 1. Table 1 reports representative frozen-trajectory contrasts, whereas the repeated-search figure reports seed-level paired summaries; the two levels of evidence are therefore read together but not pooled. In EGFR, the representative robust trajectory retained a kinase-pocket binding context while reducing dependence on a narrow mutable hotspot pattern and adding non-hotspot compensation. In HIV, the representative robust trajectory was evaluated under combo-aware optimization, so the performance annotation combines observed combo liabilities with WT feasibility.

**Table 1.**
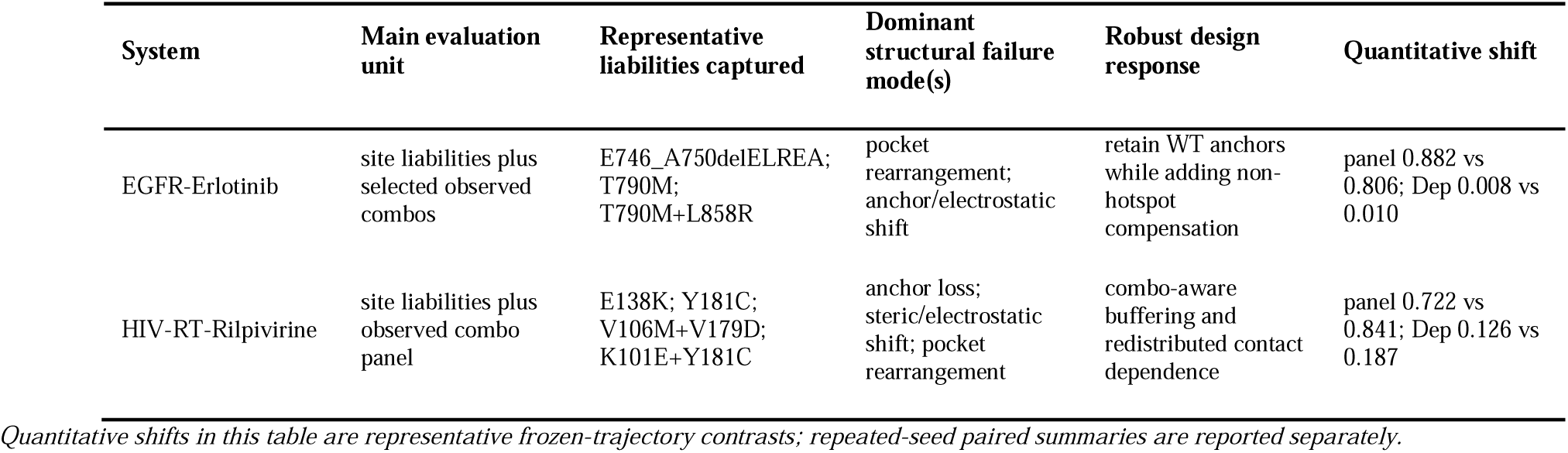
Primary-case summary of representative liabilities, dominant failure modes, and representative robust design responses.

The EGFR robust candidates illustrate the intended design response. In the representative frozen SAR contrast, robust candidates retained a kinase-pocket binding context and showed lower dependence in the Table 1 summary, whereas repeated-seed medians require a bounded interpretation. The supported conclusion is dependence reorganization with non-hotspot compensation. The most rewarded robust EGFR examples were candidates for which non-hotspot compensation and retained interaction context made the robust objective chemically legible.

The HIV robust candidates are interpreted differently because the case is combo-dense. The robust trajectory must be evaluated against site and observed-combo cores, partner-chain-aware interaction layers and non-hotspot retention. In the predefined repeated-seed summaries, robust HIV molecules can increase best robust score and preserve combo-aware scoring structure while still showing metric-level tradeoffs against naive search. This is why the manuscript describes HIV as a primary combo-aware case with its own design logic.

Contact redistribution panels show that robust-objective selections can gain or retain contacts outside the most mutable hotspot dependencies while releasing contacts that are fragile under the lead configuration (Fig. 6c). Across the two primary cases, the SAR summary highlights three positive edit-response patterns under the predefined panels: lower top-20 hotspot dependence, added non-hotspot compensation and increased scaffold-tail diversity. Non-hotspot interaction-fingerprint retention is monitored as an interaction-retention metric (Fig. 6d).

**Figure 6.**
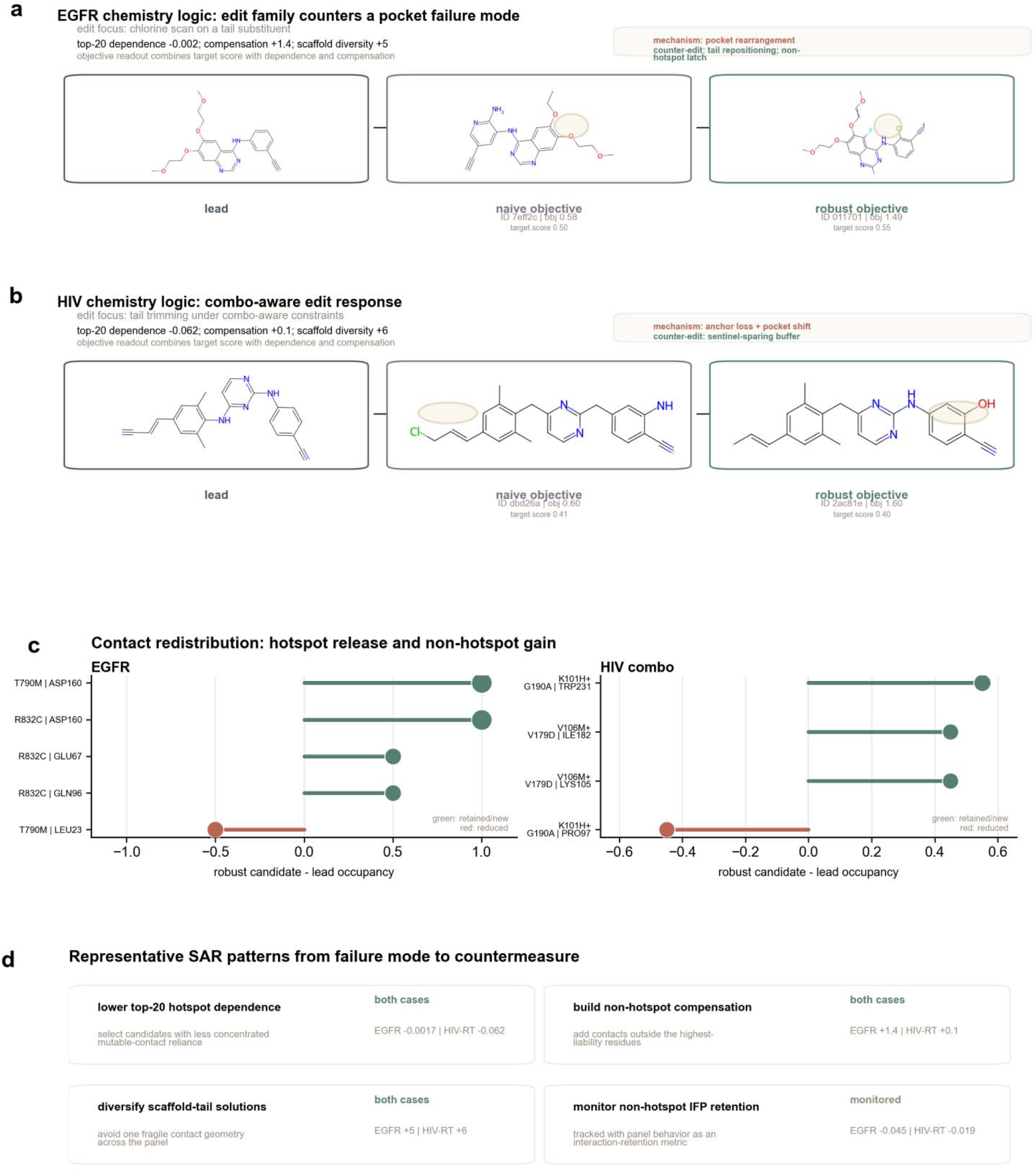
Case-specific chemistry edits reveal robust SAR from hotspot dependence to distributed robustness. (a) EGFR lead, naive-objective and robust-objective edit trajectory with failure-mode and edit-response annotation. (b) HIV combo-aware edit trajectory. (c) Contact redistribution between lead and robust-objective selections for EGFR and HIV combo settings. (d) SAR summary highlighting top-20 hotspot-dependence reduction, non-hotspot compensation, scaffold-tail diversification and monitored interaction-retention boundaries.

Together, the figure pair links objective shifts to visible chemical changes. A molecule can be evaluated as better only when the reader can see both the target-specific edit and the dependence pattern it changes; that is the logic behind the explicit Table 1 summary. The SAR conclusion is that robust counter-design can expose recurrent edit-response patterns that redistribute interaction dependence, with prospective synthesis and assay testing as the next validation step.

### Benchmark and limited physical support establish the validation envelope

These analyses test whether the closed loop is quantitatively supported beyond the two illustrative design trajectories. The short physical-support analysis used selected implicit-solvent summaries. EGFR and HIV each retained 12 successful MD-pair summaries, with 20 and 25 MM/GBSA-available summaries, respectively (Fig. 7a). These panels provide limited physical consistency support within the implicit-short envelope.

**Figure 7.**
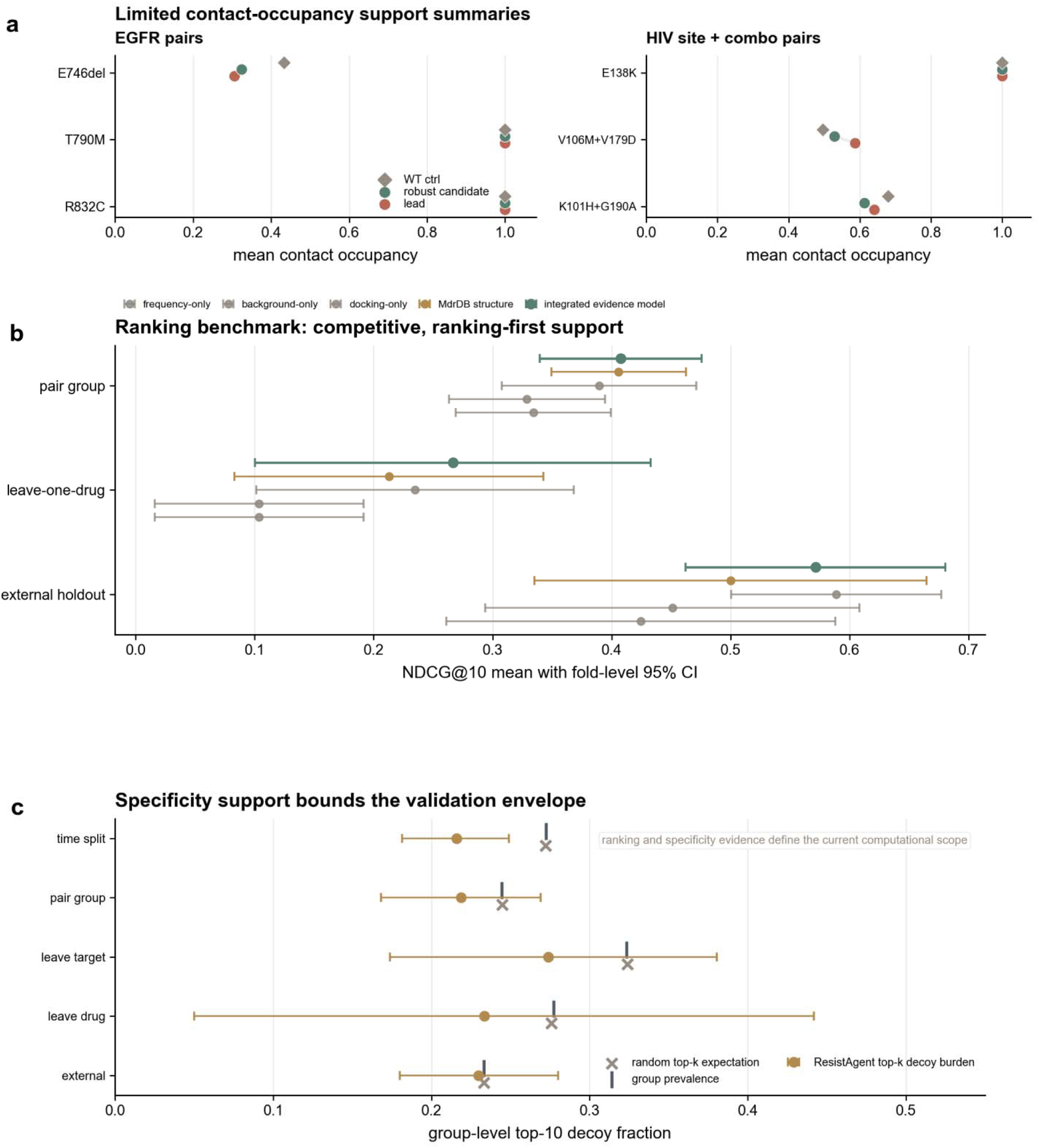
Benchmark and limited physical support define the validation envelope. (a) Selected implicit short support for EGFR and HIV site-plus-combo pairs. (b) Ranking benchmark across selected splits and baselines. (c) Decoy-specificity analysis supports benchmark interpretation.

The ranking benchmark provides ranking-first support. The selected benchmark summary includes pair-group NDCG@10 = 0.408, leave-one-drug precision@10 = 0.333, external-holdout precision@10 = 0.386 and temporal-split readiness (Fig. 7b). Decoy-specificity analysis was retained for benchmark interpretation (Fig. 7c). Detailed benchmark configuration is provided in the Supplementary Information. Competitive pair-group and precision results, together with the bounded external-holdout NDCG profile, define the benchmark as leakage-aware, ranking-first support for the selected setting.

ABL1-Nilotinib was retained only as an uncertainty-heavy boundary case, with full diagnostics reported in the Supplementary Information.

## Discussion

The central contribution of ResistAgent is conceptual and operational: mutational liabilities are moved from post hoc explanation into the objective space of lead optimization. This reframing forces three connected questions to be handled together: which mutations are plausible and damaging, how they alter interaction networks, and how counter-design can reduce dependence on mutable hotspots. The EGFR and HIV primary cases support this closed-loop narrative because they include resistance maps, mechanism diagnosis and robust SAR under predefined evidence records.

The site/combo split is not a bookkeeping detail; it changes the design problem. Once observed combinations are treated as their own liabilities, the objective no longer asks only whether a molecule can tolerate one mutation at a time. It asks whether the binding network can be reorganized so that dependence is less concentrated on the predefined mutable hotspot panel and more distributed across contacts that are less damaged by the selected liabilities. That is the substantive meaning of the phrase distributed robustness in this paper.

The robust objective also changes what counts as progress. A naive average-affinity objective can select candidates that look acceptable in aggregate while retaining fragile dependence on a small number of mutable contacts. The robust objective is designed to make lower-tail mutant behavior, panel coverage, layered interaction retention, hotspot fraction, non-hotspot compensation and synthesis realism visible at the same time. A candidate is therefore judged by whether the binding organization becomes less brittle under the predefined liability panel, not by whether one scalar affinity surrogate improves in isolation.

The validation envelope follows directly from this evidence chain. Observed or likely liabilities define computational design hypotheses under predefined panels. The LLM layer contributes evidence organization and edit rationales; deterministic engines provide physical proxies and benchmark metrics. Short implicit physical support and ranking-first benchmarks set the current evidence boundary. Within that boundary, the positive claim is that the design objective can be made resistance-aware before synthesis.

Several limitations define the next experimental steps. The chemistry edits are computationally prioritized hypotheses that require wet-lab validation. The HIV context is based on validated NNRTI holo information, and partner-chain or multichain dynamics remain targets for deeper explicit modeling before mechanistic claims are expanded. The boundary-case ABL analysis is reported in the Supplementary Information.

Together, these results establish resistance-aware lead optimization as a tractable design principle: mutational liabilities can be prioritized, mechanistically diagnosed and used to guide inhibitor edits before synthesis, while prospective assays, longer explicit-solvent ensembles and richer combo panels remain the necessary next tests.

## Materials and methods

### Data sources and case roles

The analysis combines domain-separated cancer and viral mutation evidence, WT structural models, ligand structures, interaction fingerprints and benchmark result tables. Cancer priors and viral priors are not pooled. EGFR-Erlotinib and HIV-RT-Rilpivirine are treated as the two primary cases; ABL1-Nilotinib is retained as a boundary case and its full result summary is reported in the Supplementary Information.

Structured source tables define the evidence available to each step, so the main text reports figure-level findings while detailed formulas, benchmark configuration and boundary-case diagnostics are kept in the Supplementary Information. The case roles, source-table schema and cancer/viral prior separation were fixed before manuscript assembly, and the language-model layer was not allowed to generate numerical physical scores.

### Liability construction and mapping

Mutation liabilities were constructed from appearance priors, structural-impact proxies and fitness constraints. Site liabilities and combo liabilities were explicitly separated, and observed HIV rilpivirine combinations were preserved as independently scored combo-level objects.

Counter-design panels were selected by cumulative risk mass subject to minimum and maximum panel sizes. HIV assigns a dedicated combo budget when the evaluation unit is combo-dense. The panel was a predefined site-plus-combo evaluation object fixed before counter-design. Detailed risk, weighting and scoring equations are provided in Supplementary Methods.

### Mechanism diagnosis

WT anchor networks were compared with mutant interaction fingerprints and pocket descriptors. Diagnostic features included interaction-fingerprint loss, anchor-loss fraction, pocket-volume change, local structural displacement, docking-proxy shift and MM/GBSA direction.

Mechanism labels were assigned deterministically from these structured features. MM/GBSA direction was used as a supporting calibration term. Each liability carried into counter-design retained its structured site or combo evidence record.

### Counter-design objectives and statistics

Robust and naive objectives were compared under equal search budget, including three independent paired seeds for each primary case. The robust objective emphasized lower-tail site-plus-combo panel behavior, layered interaction-fingerprint retention, hotspot-dependence reduction, non-hotspot compensation, synthesis realism and scaffold-tail diversity; the naive objective served as an average-affinity-oriented comparator.

Paired summaries report robust-minus-naive shifts for RobustScore, panel passing, raw dependence, hotspot fraction, compensation, new non-hotspot contacts and scaffold diversity. Lower dependence and hotspot fraction are preferred, so dependence deltas are interpreted in their raw direction with improvement assessed by metric preference. The main text interprets these shifts as bounded objective behavior and validation-boundary support.

### LLM role, physical support and benchmark

The language model is a bounded reasoning layer that organizes evidence, proposes constrained edit rationales and explains supported changes. Docking, MM/GBSA, FoldX-like and molecular-dynamics quantities are produced by deterministic tools, with structural quality checks and prefilters retained as binding constraints.

Selected implicit short occupancy/contact and MM/GBSA summaries provide limited physical consistency support. Ranking benchmarks use predefined leakage-aware splits and baselines; the detailed benchmark setting and weights are reported in Supplementary Table 9. Cheminformatics and ranking-model components use standard tool families including RDKit, XGBoost, scikit-learn and decoy-based benchmarking where appropriate [43–46].

## Data availability

All source data tables used to assemble the figures and tables are provided as editable source-data files with the submission package. Structured result records are retained to support reproducibility.

## Code availability

The core ResistAgent source code is publicly available at https://github.com/woshuizhaol/ResistAgent. The repository contains the analysis workflow, agent modules, configuration templates, schemas and tests.

## Acknowledgments

This work was supported by the National Natural Science Foundation of China (No. 32471313).

## CRediT authorship contribution statement

Yuxuan Wang: Conceptualization, Methodology, Software, Formal analysis, Investigation, Data curation, Validation, Visualization, Writing - original draft, Writing - review & editing. Jingyi Kang: Data curation, Validation, Literature curation, Writing - review & editing. Huizi Cui: Data curation, Validation, Literature curation, Writing - review & editing. Yu Fu: Data curation, Validation, Literature curation, Writing - review & editing. Wen Li: Supervision, Resources, Translational interpretation, Writing - review & editing. Silvio E. Perea: Supervision, Resources, Translational interpretation, Writing - review & editing. Weiwei Han: Conceptualization, Supervision, Project administration, Funding acquisition, Writing - review & editing.

## References

1. Hanahan D, Weinberg RA. Hallmarks of cancer: the next generation. Cell 2011; 144: 646–674. doi:10.1016/j.cell.2011.02.013.

2. Holohan C, Van Schaeybroeck S, Longley DB, Johnston PG. Cancer drug resistance: an evolving paradigm. Nature Reviews Cancer 2013; 13: 714–726. doi:10.1038/nrc3599.

3. Lackner MR, Wilson TR, Settleman J. Mechanisms of acquired resistance to targeted cancer therapies. Future Oncology 2012; 8: 999–1014. doi:10.2217/fon.12.86.

4. Housman G, Byler S, Heerboth S, et al.. Drug resistance in cancer: an overview. Cancers 2014; 6: 1769–1792. doi:10.3390/cancers6031769.

5. Pao W, Miller VA. Epidermal growth factor receptor mutations, small-molecule kinase inhibitors, and non-small-cell lung cancer. Journal of Clinical Oncology 2005; 23: 2556–2568. doi:10.1200/JCO.2005.07.799.

6. Lynch TJ, Bell DW, Sordella R, et al.. Activating mutations in the epidermal growth factor receptor underlying responsiveness of non-small-cell lung cancer to gefitinib. New England Journal of Medicine 2004; 350: 2129–2139. doi:10.1056/NEJMoa040938.

7. Paez JG, Janne PA, Lee JC, et al.. EGFR mutations in lung cancer: correlation with clinical response to gefitinib therapy. Science 2004; 304: 1497–1500. doi:10.1126/science.1099314.

8. Pao W, Miller V, Zakowski M, et al.. EGF receptor gene mutations are common in lung cancers from never smokers and are associated with sensitivity of tumors to gefitinib and erlotinib. Proceedings of the National Academy of Sciences 2004; 101: 13306–13311. doi:10.1073/pnas.0405220101.

9. Kobayashi S, Boggon TJ, Dayaram T, et al.. EGFR mutation and resistance of non-small-cell lung cancer to gefitinib. New England Journal of Medicine 2005; 352: 786–792. doi:10.1056/NEJMoa044238.

10. Pao W, Miller VA, Politi KA, et al.. Acquired resistance of lung adenocarcinomas to gefitinib or erlotinib is associated with a second mutation in the EGFR kinase domain. PLoS Medicine 2005; 2: e73. doi:10.1371/journal.pmed.0020073.

11. Yun CH, Boggon TJ, Li Y, et al.. Structures of lung cancer-derived EGFR mutants and inhibitor complexes: mechanism of activation and insights into differential inhibitor sensitivity. Cancer Cell 2007; 11: 217–227. doi:10.1016/j.ccr.2006.12.017.

12. Yun CH, Mengwasser KE, Toms AV, et al.. The T790M mutation in EGFR kinase causes drug resistance by increasing the affinity for ATP. Proceedings of the National Academy of Sciences 2008; 105: 2070–2075. doi:10.1073/pnas.0709662105.

13. Sequist LV, Waltman BA, Dias-Santagata D, et al.. Genotypic and histological evolution of lung cancers acquiring resistance to EGFR inhibitors. Science Translational Medicine 2011; 3: 75ra26. doi:10.1126/scitranslmed.3002003.

14. Soria JC, Ohe Y, Vansteenkiste J, et al.. Osimertinib in untreated EGFR-mutated advanced non-small-cell lung cancer. New England Journal of Medicine 2018; 378: 113–125. doi:10.1056/NEJMoa1713137.

15. Rhee SY, Gonzales MJ, Kantor R, Betts BJ, Ravela J, Shafer RW. Human immunodeficiency virus reverse transcriptase and protease sequence database. Nucleic Acids Research 2003; 31: 298–303. doi:10.1093/nar/gkg100.

16. Bennett DE, Camacho RJ, Otelea D, et al.. Drug resistance mutations for surveillance of transmitted HIV-1 drug-resistance: 2009 update. PLoS ONE 2009; 4: e4724. doi:10.1371/journal.pone.0004724.

17. Wensing AM, Calvez V, Gunthard HF, et al.. 2017 update of the drug resistance mutations in HIV-1. Topics in Antiviral Medicine 2017; 24: 132–133.

18. Shafer RW. Rationale and uses of a public HIV drug-resistance database. Journal of Infectious Diseases 2006; 194 Suppl 1: S51–S58. doi:10.1086/505356.

19. De Clercq E. Non-nucleoside reverse transcriptase inhibitors (NNRTIs): past, present, and future. Chemistry & Biodiversity 2004; 1: 44–64. doi:10.1002/cbdv.200490012.

20. Das K, Bauman JD, Clark AD Jr, et al.. High-resolution structures of HIV-1 reverse transcriptase/TMC278 complexes: strategic flexibility explains potency against resistance mutations. Proceedings of the National Academy of Sciences 2008; 105: 1466–1471. doi:10.1073/pnas.0711209105.

21. Ren J, Nichols C, Bird LE, et al.. Structural basis for the resilience of efavirenz to drug resistance mutations in HIV-1 reverse transcriptase. Structure 2000; 8: 1089–1094. doi:10.1016/S0969-2126(00)00513-X.

22. Hsiou Y, Ding J, Das K, et al.. Structural mechanisms of drug resistance for mutations at codons 181 and 188 in HIV-1 reverse transcriptase and the improved resilience of second generation non-nucleoside inhibitors. Journal of Molecular Biology 2001; 309: 437–445. doi:10.1006/jmbi.2001.4988.

23. Menendez-Arias L. Molecular basis of human immunodeficiency virus type 1 drug resistance: overview and recent developments. Antiviral Research 2013; 98: 93–120. doi:10.1016/j.antiviral.2013.01.007.

24. Clavel F, Hance AJ. HIV drug resistance. New England Journal of Medicine 2004; 350: 1023–1035. doi:10.1056/NEJMra025195.

25. Deeks SG. Nonnucleoside reverse transcriptase inhibitor resistance. Journal of Acquired Immune Deficiency Syndromes 2001; 26 Suppl 1: S25–S33. doi:10.1097/00126334-200101011-00005.

26. Berman HM, Westbrook J, Feng Z, et al.. The Protein Data Bank. Nucleic Acids Research 2000; 28: 235–242. doi:10.1093/nar/28.1.235.

27. Burley SK, Bhikadiya C, Bi C, et al.. RCSB Protein Data Bank: powerful new tools for exploring 3D structures of biological macromolecules. Nucleic Acids Research 2021; 49: D437–D451. doi:10.1093/nar/gkaa1038.

28. The UniProt Consortium. UniProt: the Universal Protein Knowledgebase in 2023. Nucleic Acids Research 2023; 51: D523–D531. doi:10.1093/nar/gkac1052.

29. Tate JG, Bamford S, Jubb HC, et al.. COSMIC: the Catalogue Of Somatic Mutations In Cancer. Nucleic Acids Research 2019; 47: D941–D947. doi:10.1093/nar/gky1015.

30. Sondka Z, Bamford S, Cole CG, Ward SA, Dunham I, Forbes SA. The COSMIC Cancer Gene Census: describing genetic dysfunction across all human cancers. Nature Reviews Cancer 2018; 18: 696–705. doi:10.1038/s41568-018-0060-1.

31. Meyers RM, Bryan JG, McFarland JM, et al.. Computational correction of copy number effect improves specificity of CRISPR-Cas9 essentiality screens in cancer cells. Nature Genetics 2017; 49: 1779–1784. doi:10.1038/ng.3984.

32. Tsherniak A, Vazquez F, Montgomery PG, et al.. Defining a cancer dependency map. Cell 2017; 170: 564–576.e16. doi:10.1016/j.cell.2017.06.010.

33. Wishart DS, Feunang YD, Guo AC, et al.. DrugBank 5.0: a major update to the DrugBank database for 2018. Nucleic Acids Research 2018; 46: D1074–D1082. doi:10.1093/nar/gkx1037.

34. Mendez D, Gaulton A, Bento AP, et al.. ChEMBL: towards direct deposition of bioassay data. Nucleic Acids Research 2019; 47: D930–D940. doi:10.1093/nar/gky1075.

35. Kim S, Chen J, Cheng T, et al.. PubChem 2023 update. Nucleic Acids Research 2023; 51: D1373–D1380. doi:10.1093/nar/gkac956.

36. Gilson MK, Liu T, Baitaluk M, Nicola G, Hwang L, Chong J. BindingDB in 2015: a public database for medicinal chemistry, computational chemistry and systems pharmacology. Nucleic Acids Research 2016; 44: D1045–D1053. doi:10.1093/nar/gkv1072.

37. Wang R, Fang X, Lu Y, Wang S. The PDBbind database: collection of binding affinities for protein-ligand complexes with known three-dimensional structures. Journal of Medicinal Chemistry 2004; 47: 2977–2980. doi:10.1021/jm030580l.

38. Trott O, Olson AJ. AutoDock Vina: improving the speed and accuracy of docking with a new scoring function, efficient optimization, and multithreading. Journal of Computational Chemistry 2010; 31: 455–461. doi:10.1002/jcc.21334.

39. Eberhardt J, Santos-Martins D, Tillack AF, Forli S. AutoDock Vina 1.2.0: new docking methods, expanded force field, and Python bindings. Journal of Chemical Information and Modeling 2021; 61: 3891–3898. doi:10.1021/acs.jcim.1c00203.

40. Salentin S, Schreiber S, Haupt VJ, Adasme MF, Schroeder M. PLIP: fully automated protein-ligand interaction profiler. Nucleic Acids Research 2015; 43: W443–W447. doi:10.1093/nar/gkv315.

41. Kollman PA, Massova I, Reyes C, et al.. Calculating structures and free energies of complex molecules: combining molecular mechanics and continuum models. Accounts of Chemical Research 2000; 33: 889–897. doi:10.1021/ar000033j.

42. Homeyer N, Gohlke H. Free energy calculations by the molecular mechanics Poisson-Boltzmann surface area method. Molecular Informatics 2012; 31: 114–122. doi:10.1002/minf.201100135.

43. Landrum G. RDKit: open-source cheminformatics software. Zenodo 2024. doi:10.5281/zenodo.591637.

44. Chen T, Guestrin C. XGBoost: a scalable tree boosting system. Proceedings of the 22nd ACM SIGKDD International Conference on Knowledge Discovery and Data Mining 2016: 785–794. doi:10.1145/2939672.2939785.

45. Pedregosa F, Varoquaux G, Gramfort A, et al.. Scikit-learn: machine learning in Python. Journal of Machine Learning Research 2011; 12: 2825–2830.

46. Mysinger MM, Carchia M, Irwin JJ, Shoichet BK. Directory of Useful Decoys, Enhanced (DUD-E): better ligands and decoys for better benchmarking. Journal of Medicinal Chemistry 2012; 55: 6582–6594. doi:10.1021/jm300687e.

