## Supplementary Information for "From hotspot dependence to distributed robustness in resistance-aware lead optimization"

This Supplementary Information contains Supplementary Methods, Supplementary Notes, Supplementary Figures, Supplementary Tables and Supplementary References. Large data tables are supplied as an editable Excel workbook, with figures reserved for graphical evidence.

### Contents

Supplementary Methods

Supplementary Notes 1-6

Supplementary Figures 1-14

Supplementary Tables 1-11

Supplementary References

### Supplementary Methods

The supplementary methods provide additional methodological detail supporting the main paper and the accompanying source-data workbook.

#### Data and prior construction

Cancer priors and viral priors were maintained separately. Blind benchmark priors excluded same-drug leakage where required for ranking evaluation. Source database roles, evidence tiers and guardrails are summarized in Supplementary Table 1.

#### HIV site-plus-combo handling

HIV-RT-Rilpivirine was treated as a combo-dense case. Observed rilpivirine mutation combinations were carried forward as combo-level liabilities and evaluated separately from site-level liabilities.

#### Structural quality control and WT anchors

WT complexes, redocking attempts, pocket context and anchor residues were used to define the structural coordinate system. HIV main-text structural interpretation uses validated NNRTI holo context as the reference structural basis.

#### Counter-design diagnostics

Search diagnostics include valid candidate rate, panel coverage, prefilter behavior and objective ablation. Robust candidates are interpreted through dependence redistribution, non-hotspot compensation and affinity behavior together.

#### Supplementary physical support

Selected implicit short contact/occupancy and MM/GBSA summaries provide limited physical consistency evidence and are not used as a long-timescale explicit-solvent validation package.

#### Benchmark and specificity

The ranking benchmark provides competitive but bounded support. Full split and baseline metrics are provided in Supplementary Figure 13 and Supplementary Table 8. Group-level decoy specificity is retained to interpret benchmark behavior and not used as a broad model-dominance claim.

#### Supplementary Methods: mathematical definitions

The main text reports the design logic and primary results. The explicit mathematical definitions used for priors, liability scoring, panel weighting, robust objectives, interaction retention, dependence, compensation, reward and evidence-confidence weighting are provided here.

**Appearance prior**

P_bg denotes baseline or population-level occurrence, and P_sel denotes drug-selected or same-drug evidence when allowed for production case analysis.

$$P_{\mathrm{app}}(i)=1-\left( 1-P_{\mathrm{bg}}(i) \right)\left( 1-P_{\mathrm{sel}}(i) \right)$$

**Raw explanatory risk**

F_i denotes the fitness constraint term.

$$R_{\mathrm{raw}}(i)=P_{\mathrm{app}}(i) I_{i} F_{i}$$

**Impact proxy**

Delta D_i and Delta Q_i denote docking-impact and interaction-fingerprint impact proxies with group-wise rank normalization.

$$I_{i}=\mathrm{ranknorm}_{g}(\Delta D_{i})+0.5\mathrm{ranknorm}_{g}(\Delta Q_{i})$$

**Panel weight**

r_i is the liability score and tau is the panel-weight temperature.

$$w_{i}=W_{s}\frac{exp(log(max(r_{i},\epsilon)+\epsilon)/\tau)}{\sum_{j} \exp(log(max(r_{j},\epsilon)+\epsilon)/\tau)}$$

**Candidate-target binding score**

E is the docking-affinity proxy and sigma is the binding-score scale.

$$S(c,t)=\frac{1}{1+exp[-(E_{lead,t}-E_{c,t})/\sigma]}$$

**Uncertainty-heavy target score**

KIFP denotes layered interaction-fingerprint retention and L_anchor is the anchor-loss term.

$$S_{u}=0.4S_{b}+0.4KIFP+0.2(1-L_{\mathrm{anchor}})$$

**Weighted lower-tail robust core**

CVaR is computed over the selected mutation panel with liability weights.

$$R_{\mathrm{core}}(c)=\mathrm{CVaR}_{q}(\{S(c,t)\},\{w_{t}\}), q=0.20$$

**WT-constrained robust score**

The robust score is constrained by both WT feasibility and mutant-tail behavior.

$$R_{\mathrm{rob}}(c)=min(S(c,WT),R_{\mathrm{core}}(c))$$

**Naive mean comparator**

The naive comparator averages WT and weighted panel behavior.

$$R_{\mathrm{naive}}(c)=\frac{S(c,WT)+\mathrm{wmean}_{t}S(c,t)}{2}$$

**Layered interaction-fingerprint retention**

Q_l denotes a layer-specific interaction fingerprint and a denotes the anchor reference.

$$KIFP(c)=\sum_{l} a_{l}cos(Q_{l}(c),Q_{l}(a))$$

**Hotspot dependence**

h(c) is hotspot contact mass divided by total pocket contact mass.

$$D_{\mathrm{dep}}(c)=0.5h(c)+0.5(1-D_{\mathrm{all}}(c))$$

**Non-hotspot compensation**

N_nh denotes non-hotspot contact support.

$$G_{\mathrm{comp}}(c)=max(0,N_{\mathrm{nh}}(c)-N_{\mathrm{nh}}(lead))$$

**Robust reward**

The reward combines WT feasibility, site and combo cores, alternative anchors, non-hotspot support and penalties.

$$R_{v2}(c)=0.8S_{\mathrm{WT}}+1.0R_{\mathrm{site}}+1.2I_{\mathrm{combo}}R_{\mathrm{combo}}+0.5A_{\mathrm{alt}}+0.35N_{\mathrm{nh}}-P_{\mathrm{pen}}(c)$$

**Reward penalties**

The penalty term combines hotspot fraction, evidence uncertainty and synthesis penalty.

$$P_{\mathrm{pen}}(c)=0.25H_{\mathrm{frac}}+0.25U_{\mathrm{evid}}+0.25P_{\mathrm{synth}}$$

**Evidence-confidence weighting**

Confidence combines calibration, sample support, target coverage and missingness/uncertainty terms.

$$T=0.35\rho_{P}+0.15\rho_{S}+0.15min(n_{\mathrm{train}}/20,1)+0.15min(n_{\mathrm{targets}}/10,1)+0.10(1-r_{u})+0.10(1-r_{m})$$

The numerical settings needed to read these equations and repeated-search summaries are collected in Supplementary Table 11 and the editable supplementary workbook.

### Supplementary Notes

#### Supplementary Note 1. Role of the LLM layer

The LLM layer organizes evidence, generates constrained explanations and proposes supported edit templates. Docking, MM/GBSA, FoldX-like and molecular-dynamics quantities are produced by deterministic tools, and the design decisions remain evidence-constrained.

#### Supplementary Note 2. Why site and combo liabilities are separated

Site-level liabilities answer which individual mutable positions should be considered, whereas combo-level liabilities preserve observed multi-mutation patterns. HIV-RT-Rilpivirine requires this distinction because the observed combo panel carries mechanistic and evolutionary information not represented by independent site scores.

#### Supplementary Note 3. How short physical support is interpreted

The physical-support analysis uses implicit short summaries and selected pair-level evidence. These data support plausibility and consistency but do not replace explicit-solvent validation or experimental binding assays.

#### Supplementary Note 4. Why ABL is a boundary case

ABL1-Nilotinib has weaker dock-versus-MM/GBSA calibration and uncertainty-heavy evidence-confidence behavior. It is therefore used to characterize boundary-case failure modes, not to support the same level of primary evidence as EGFR and HIV.

#### Supplementary Note 5. Evidence boundary of the ranking benchmark

The benchmark provides competitive ranking-first support under leakage-controlled split and prior settings. It should not be interpreted as superiority across all possible baselines, splits, metrics or specificity definitions.

#### Supplementary Note 6. ABL result relocation

The full ABL1-Nilotinib boundary-case result summary is kept in Supplementary Table 10. This preserves ABL as a boundary case while keeping the main narrative centered on EGFR-Erlotinib and HIV-RT-Rilpivirine.

### Supplementary Figures

**Supplementary Figure 1. Master-table evaluation-unit landscape and case-role distribution supporting the main design-loop overview.**

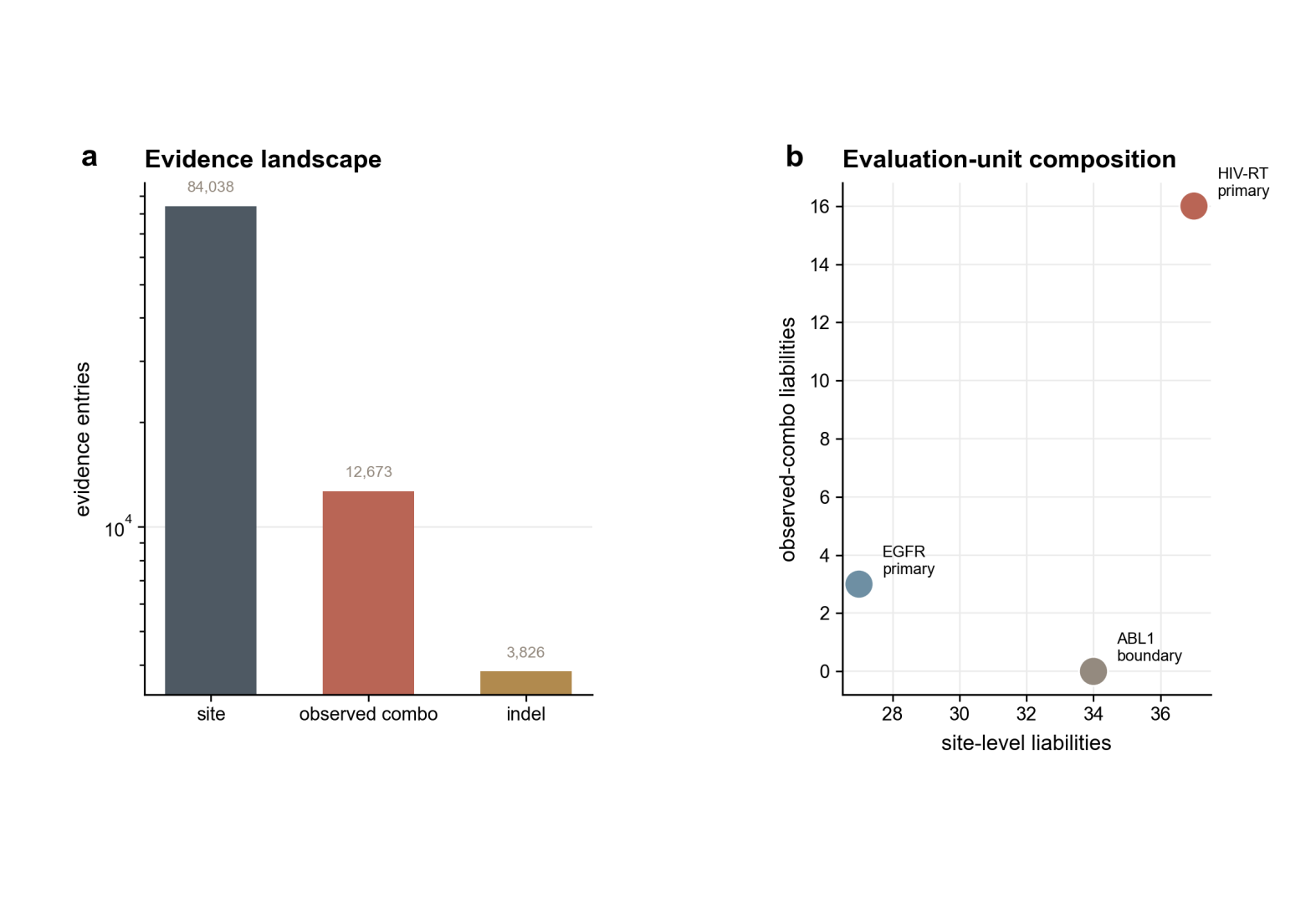

**Supplementary Figure 2. Prior construction and blind-prior control, including domain-separated viral and cancer prior handling.**

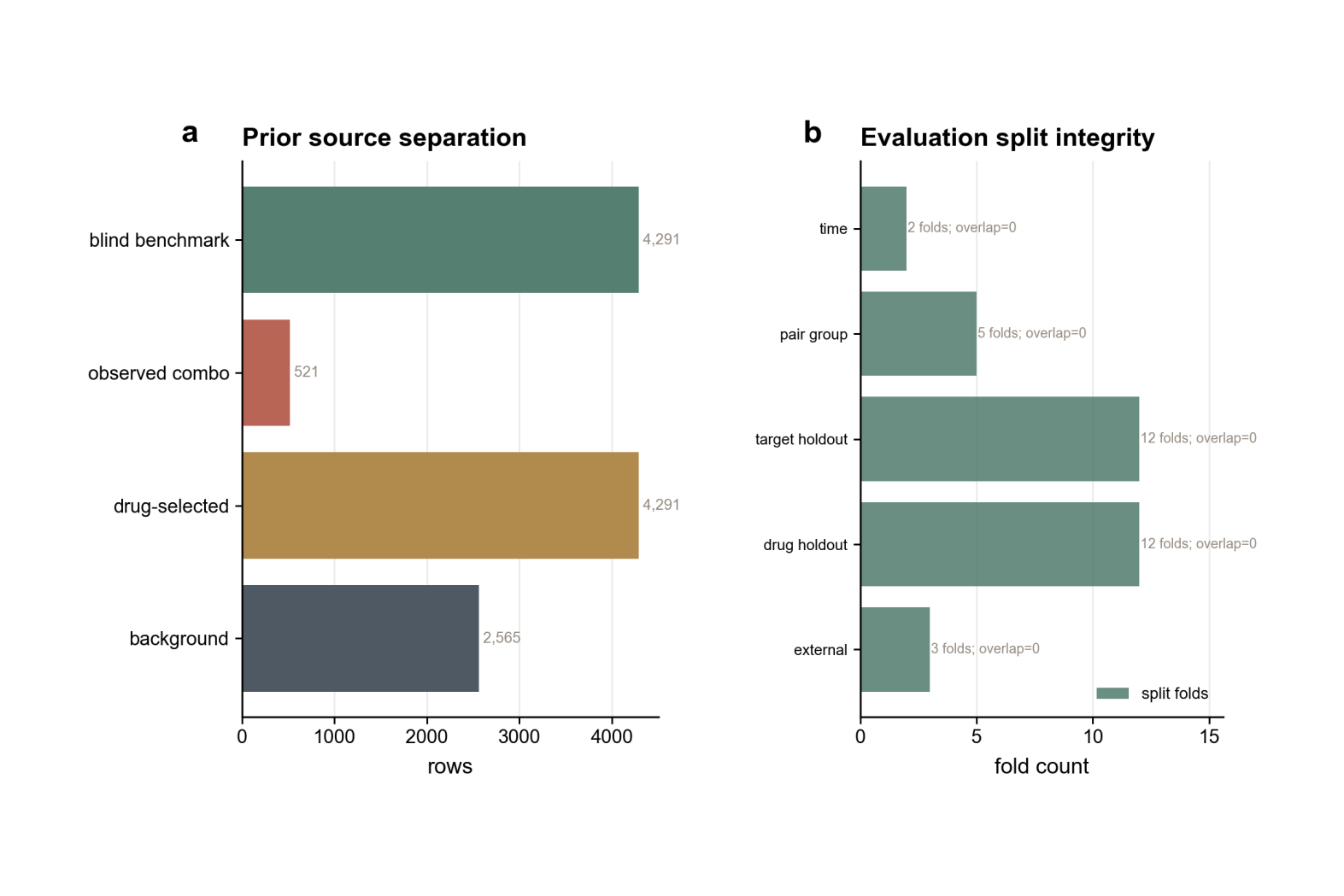

**Supplementary Figure 3. HIV RT domain extraction, NNRTI holo support and pocket-context summary.**

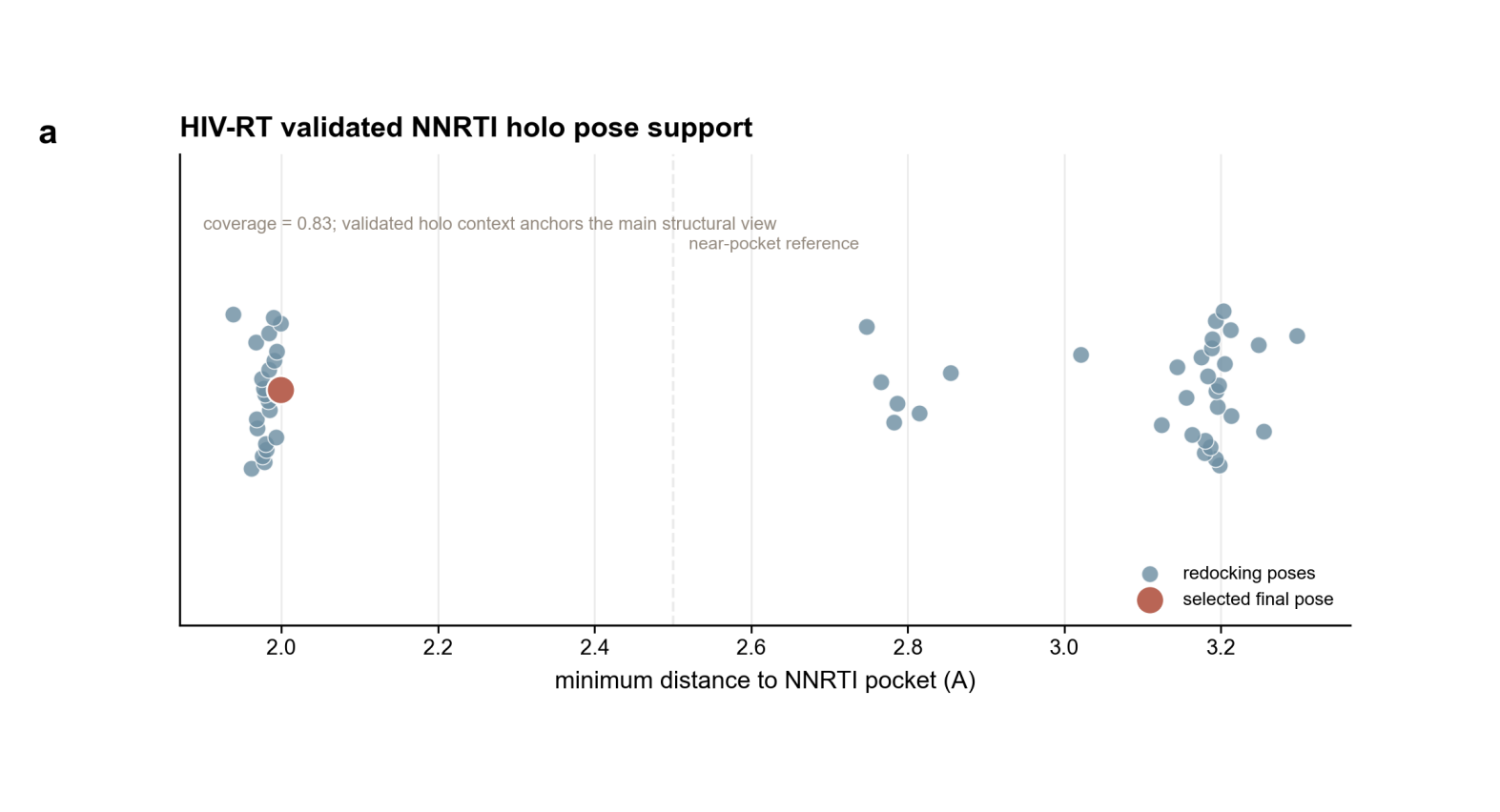

**Supplementary Figure 4. Residue numbering, component mapping and combo-entry coverage summary.**

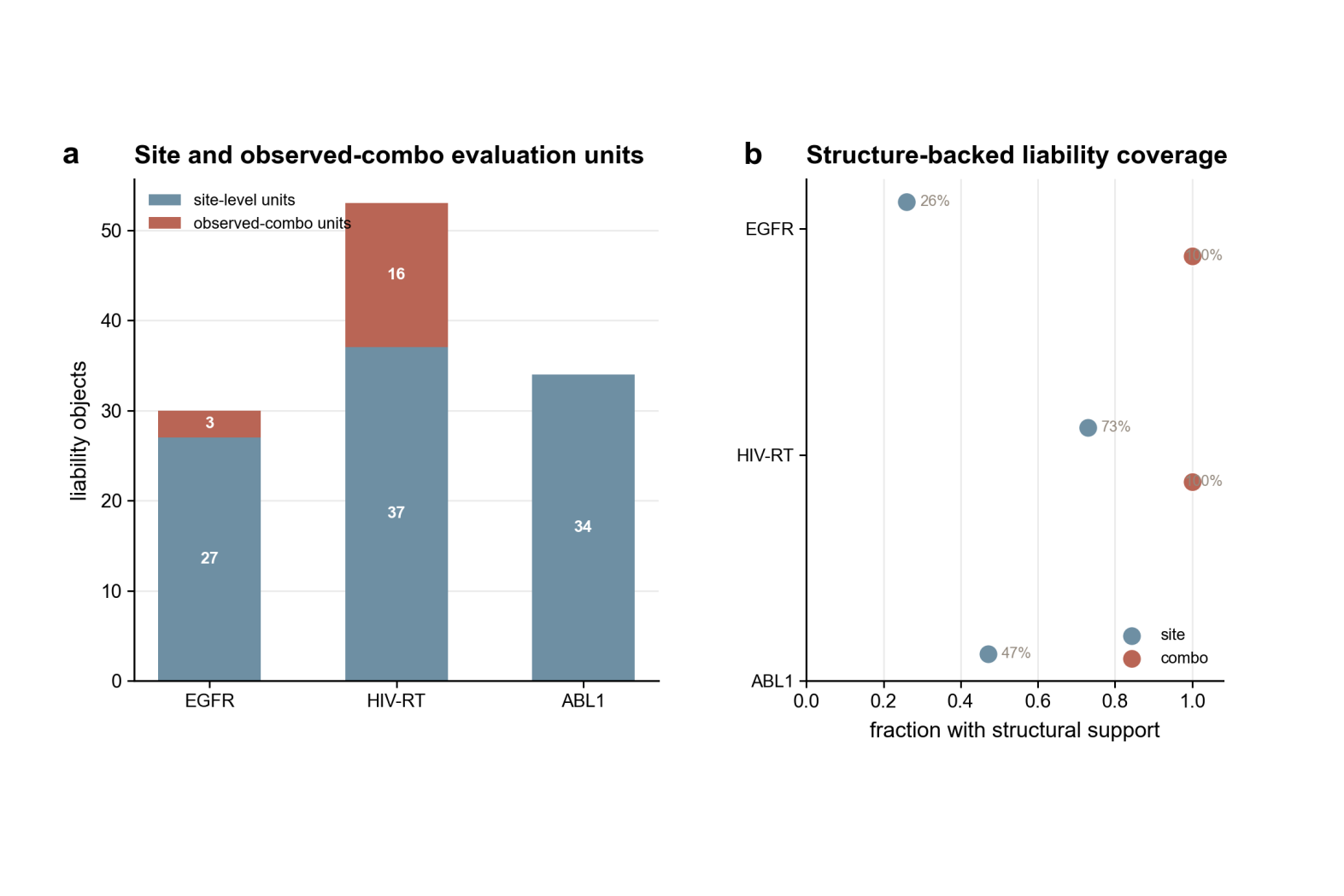

**Supplementary Figure 5. WT pose, anchor-pose and docking-context validation summaries.**

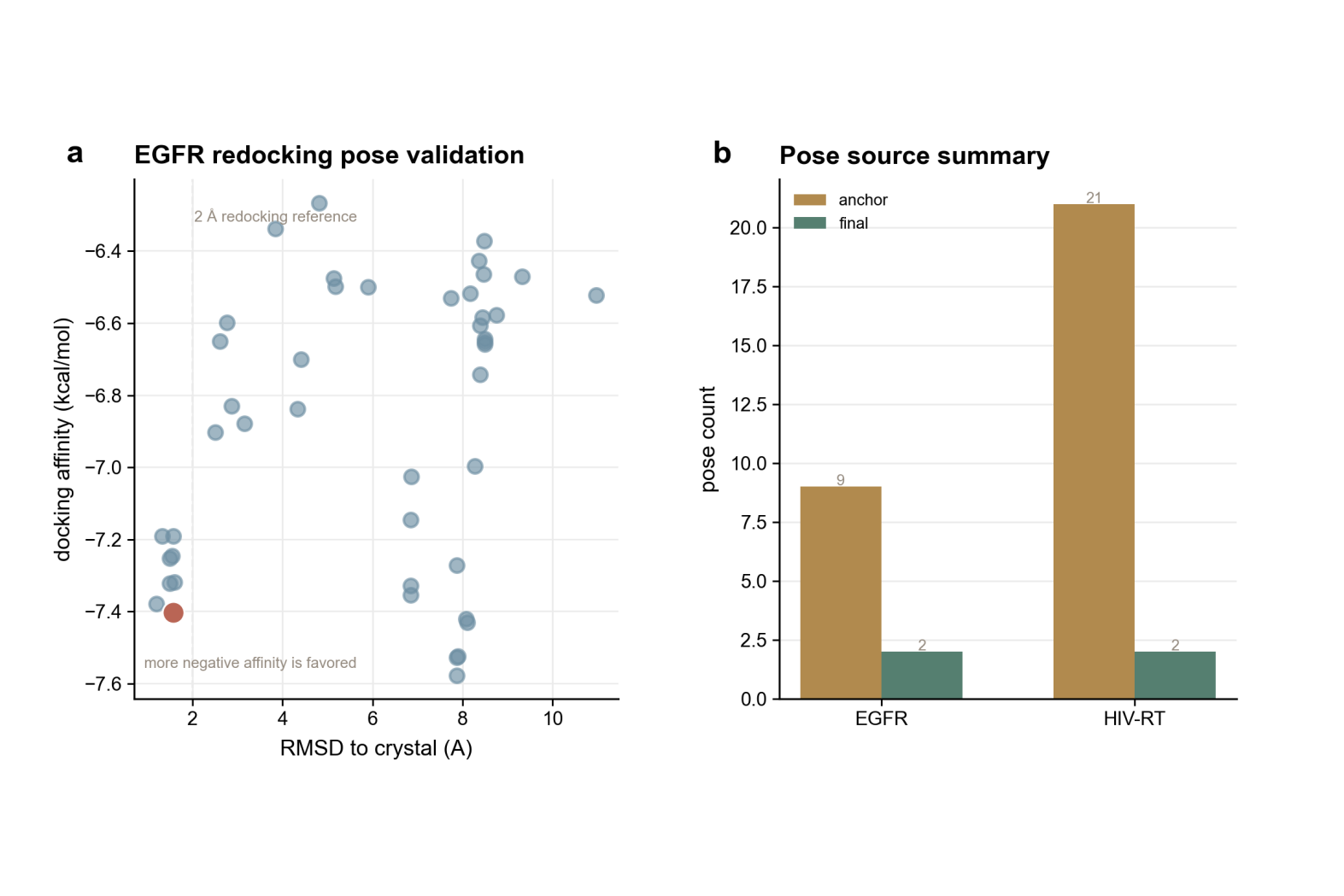

**Supplementary Figure 6. Full-resolution EGFR, HIV and ABL residue maps supporting the condensed Figure 3 panels.**

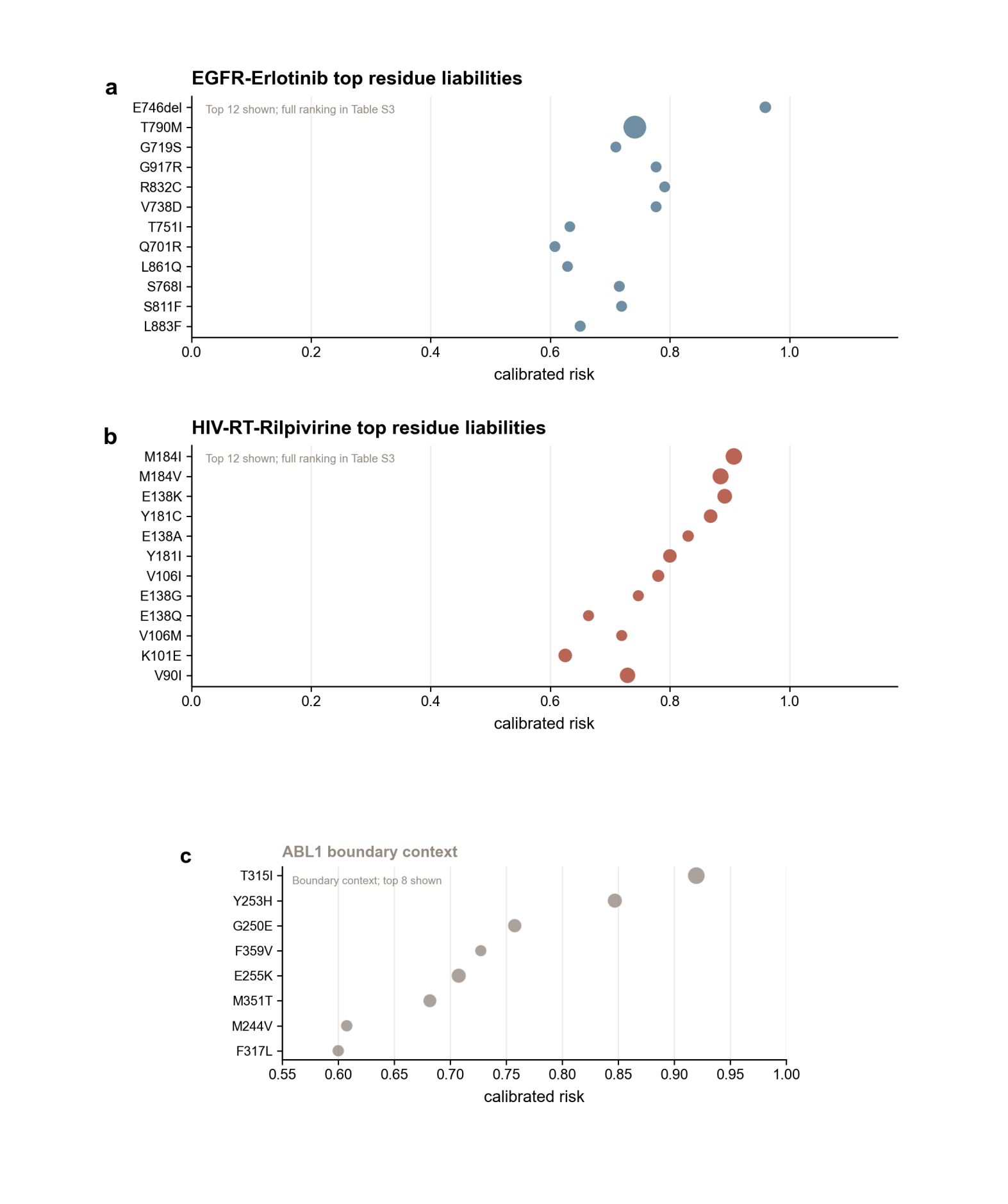

**Supplementary Figure 7. Full combo rankings and additional observed-combo exemplars for the HIV case.**

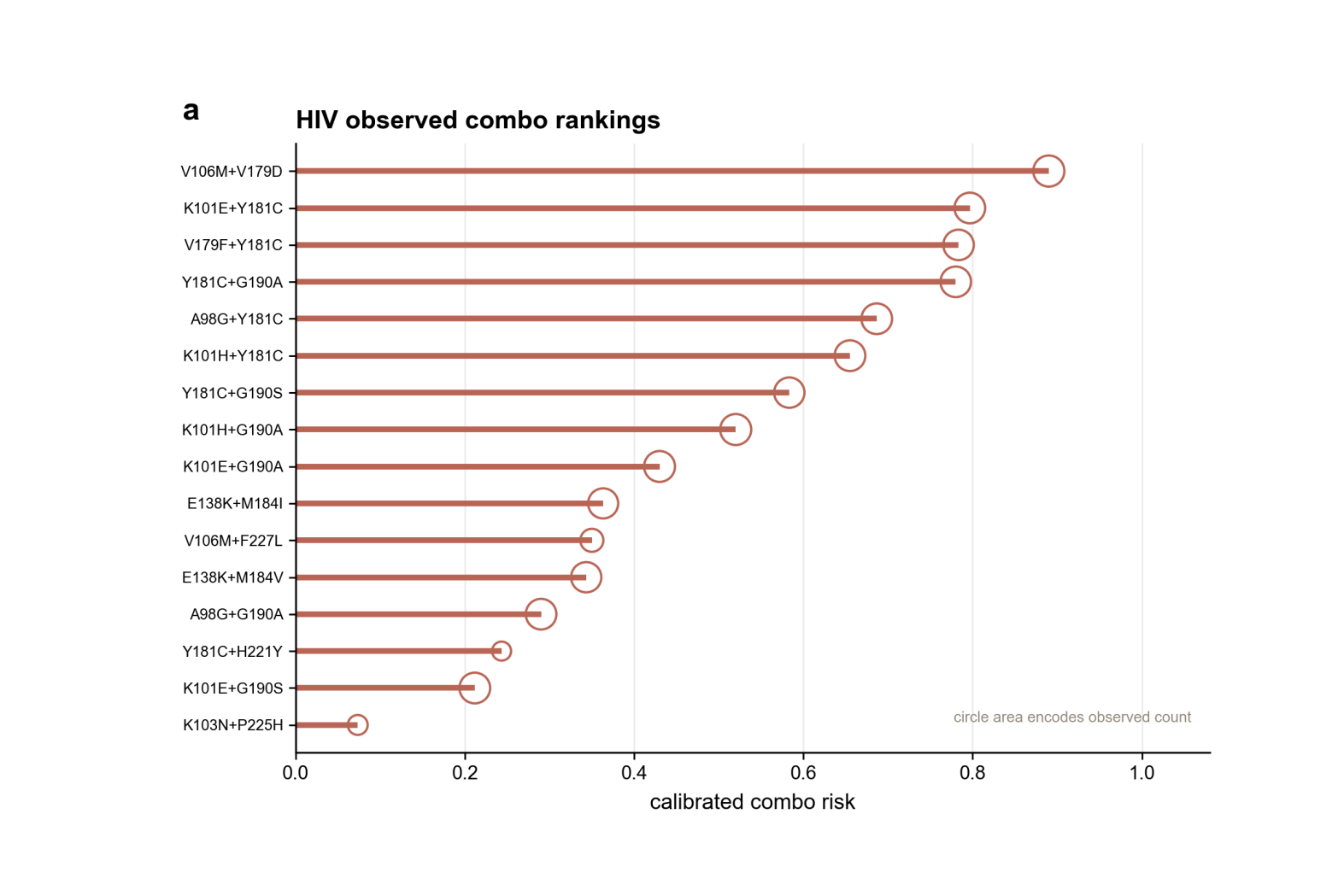

**Supplementary Figure 8. Extended mechanism and calibration analysis, including ABL boundary-case calibration.**

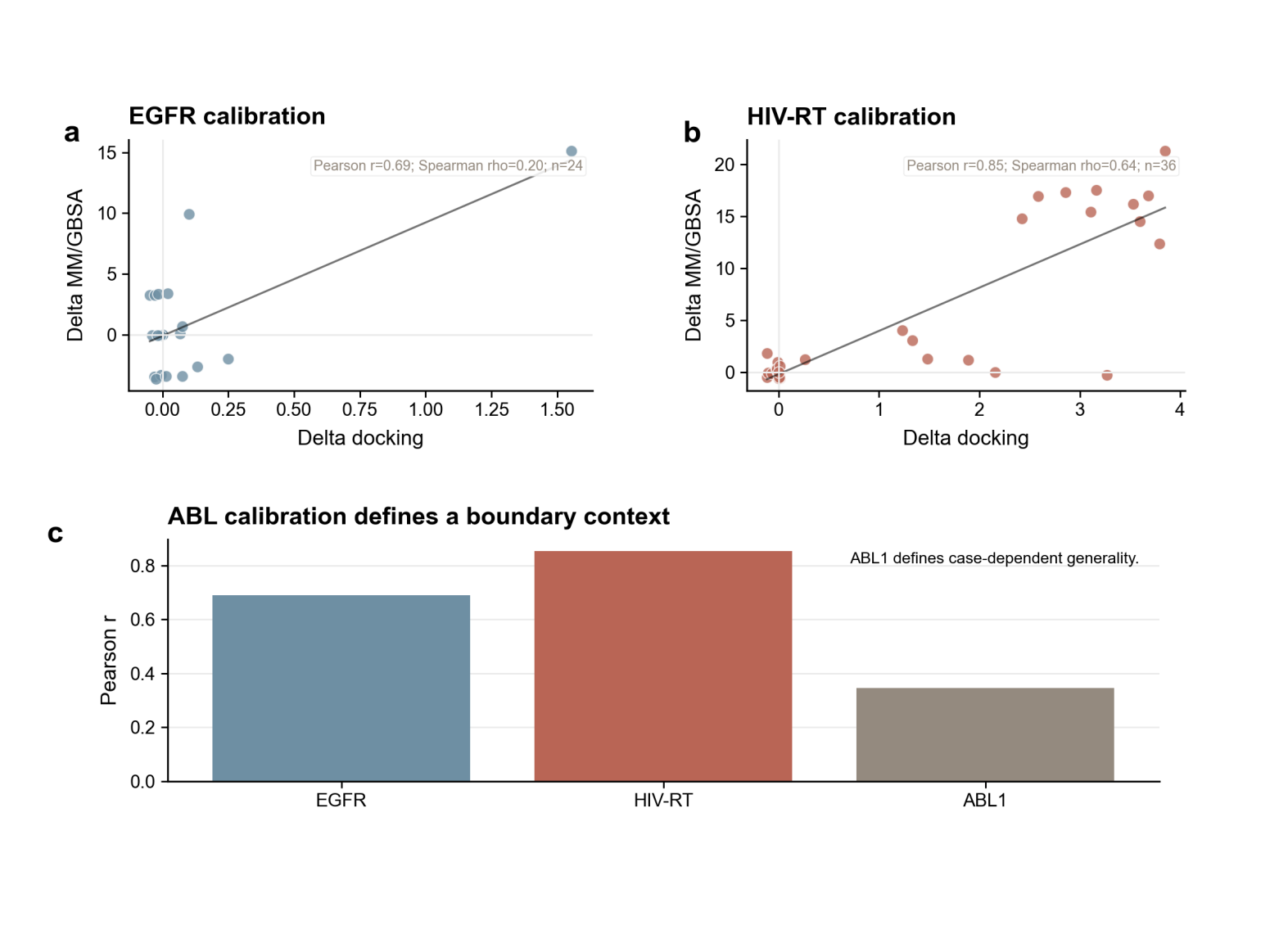

**Supplementary Figure 9. Counter-design search trajectory, candidate-retention and design-constraint summaries.**

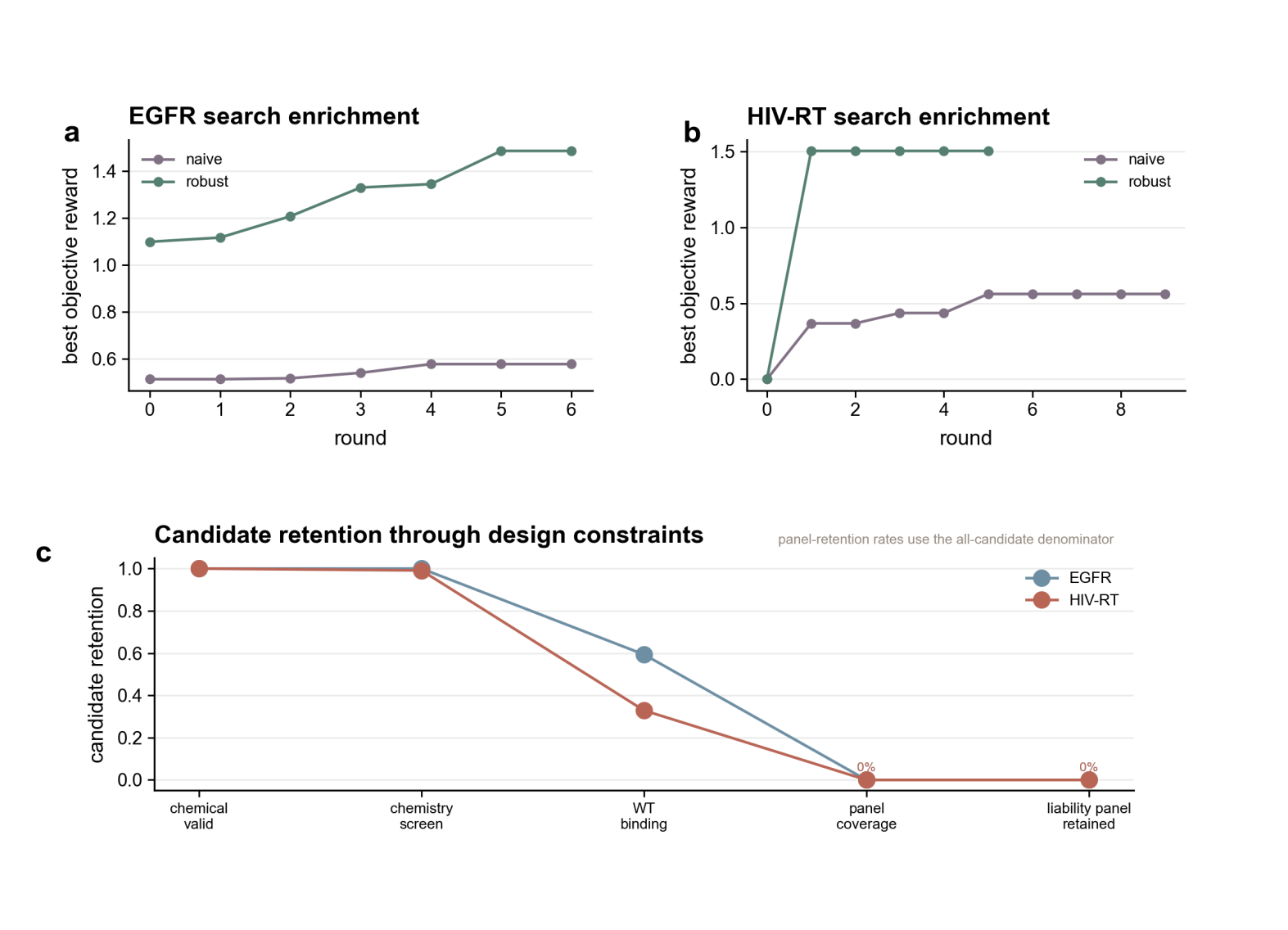

**Supplementary Figure 10. Objective ablations for robust and average-affinity-oriented search settings, including HIV combo-aware sensitivity.**

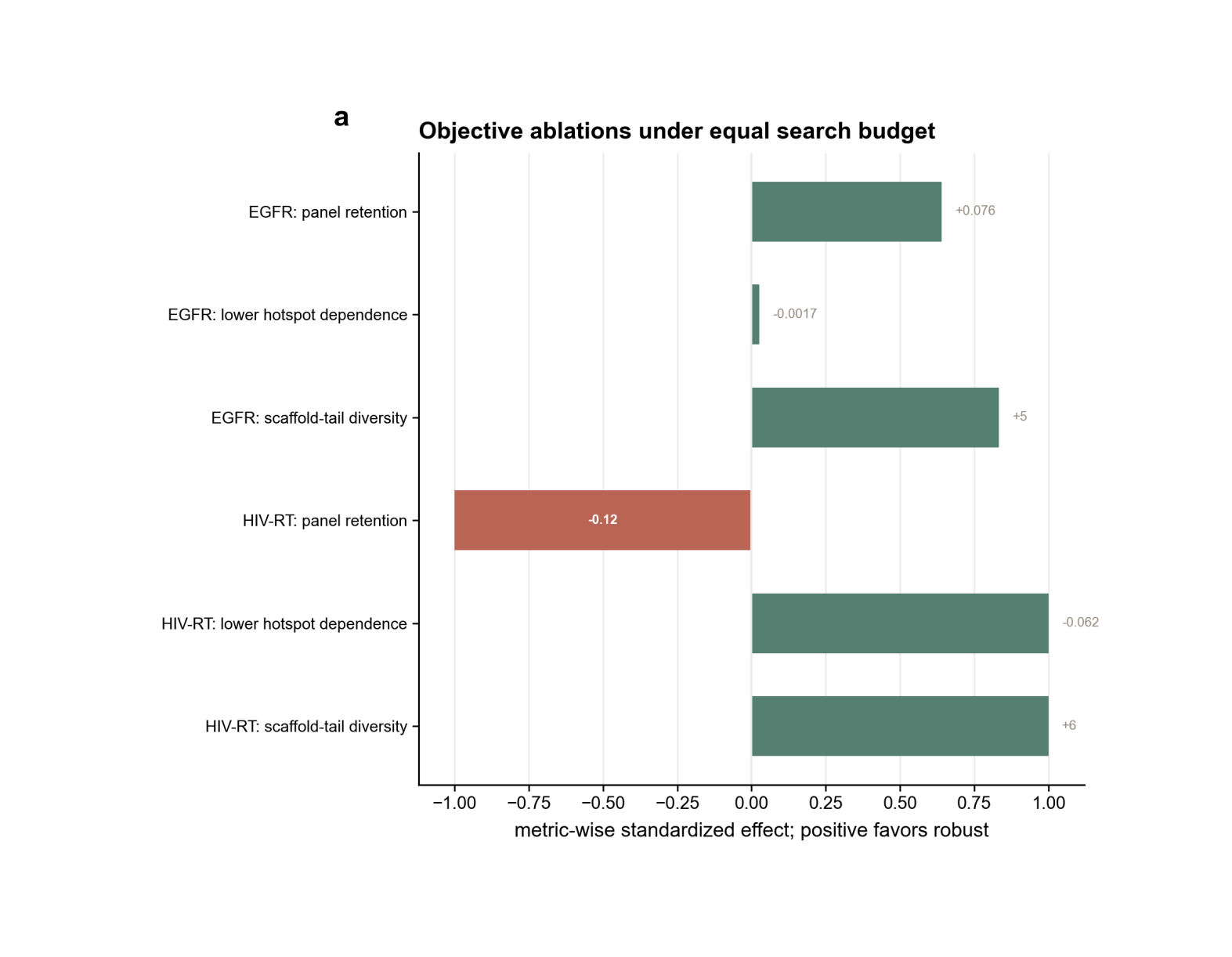

**Supplementary Figure 11. ABL boundary-case and uncertainty-heavy evidence summary supporting its non-primary assignment.**

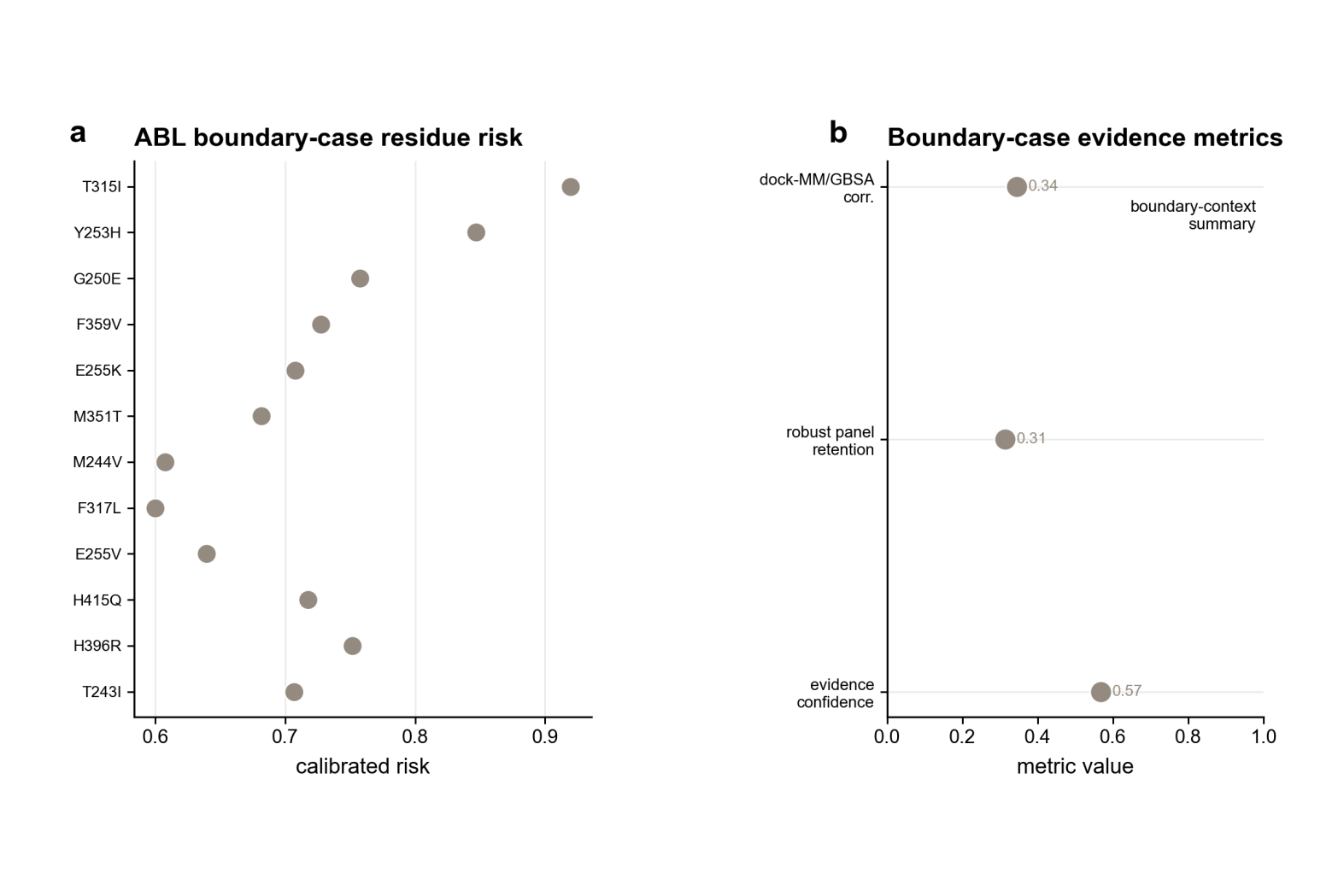

**Supplementary Figure 12. Pair-level occupancy and MM/GBSA consistency for supplementary implicit short physical support.**

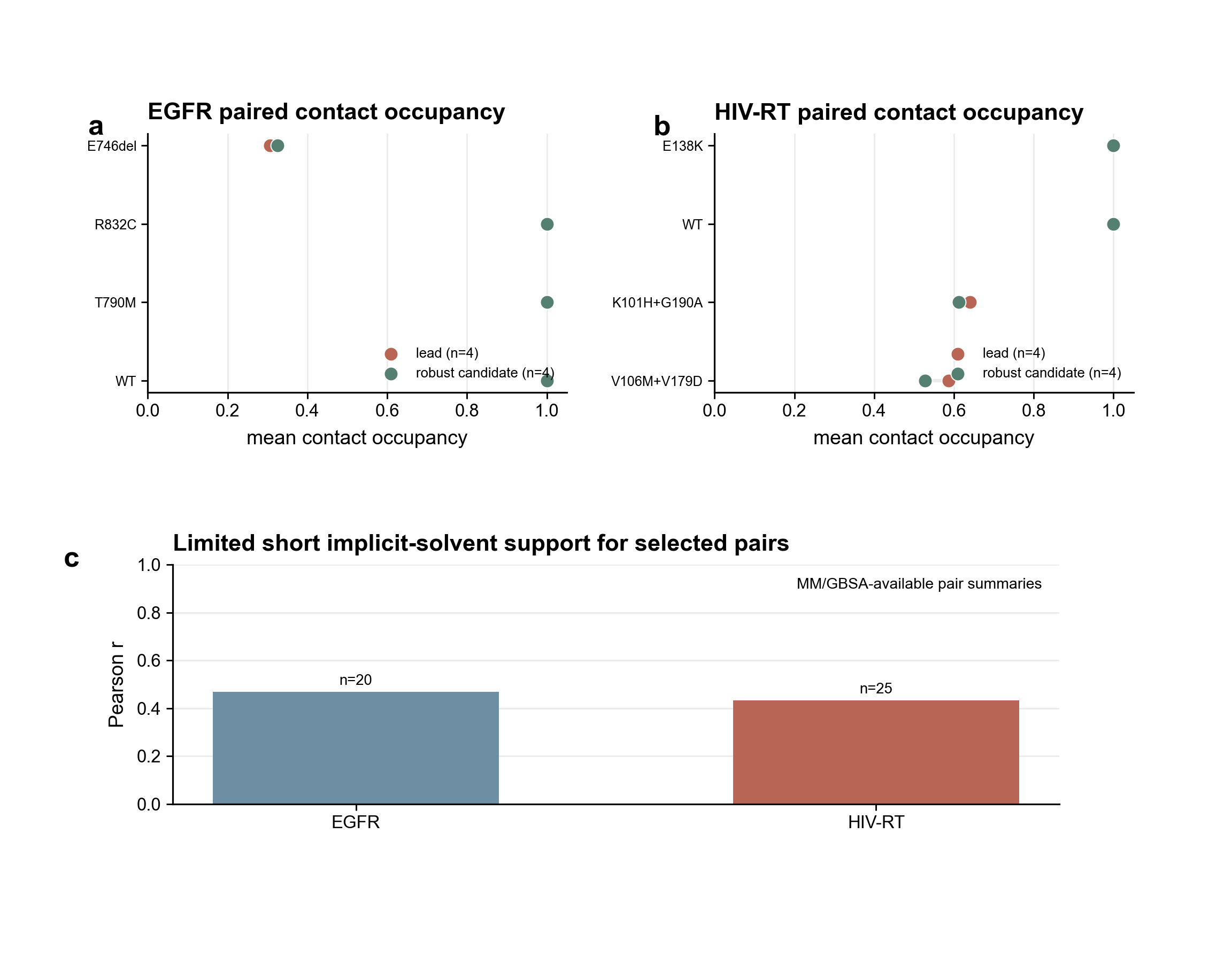

**Supplementary Figure 13. Full ranking benchmark across splits, baselines and subgroups.**

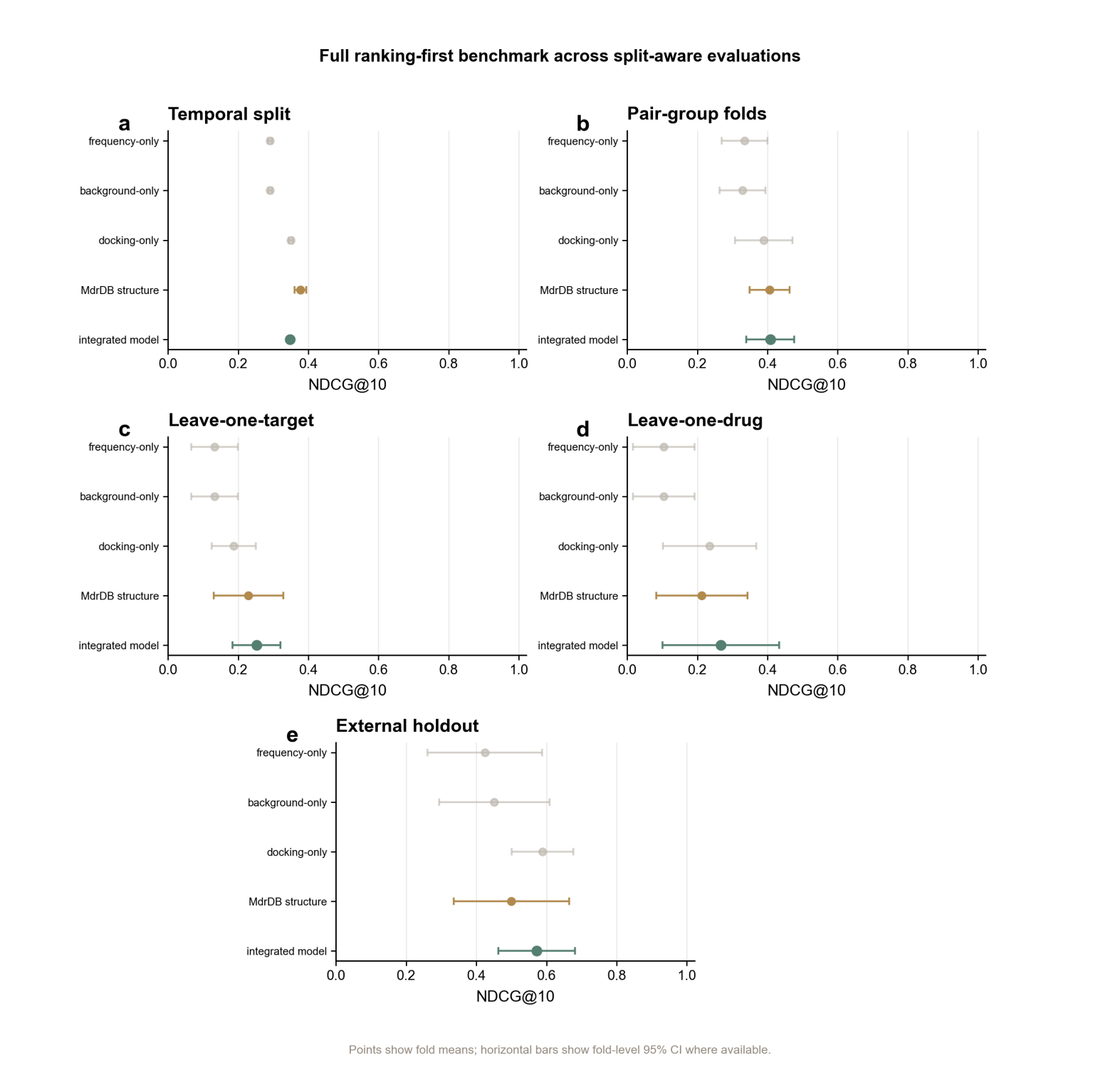

**Supplementary Figure 14. Decoy-specificity and prior-control summary; decoy specificity is used to interpret benchmark behavior.**

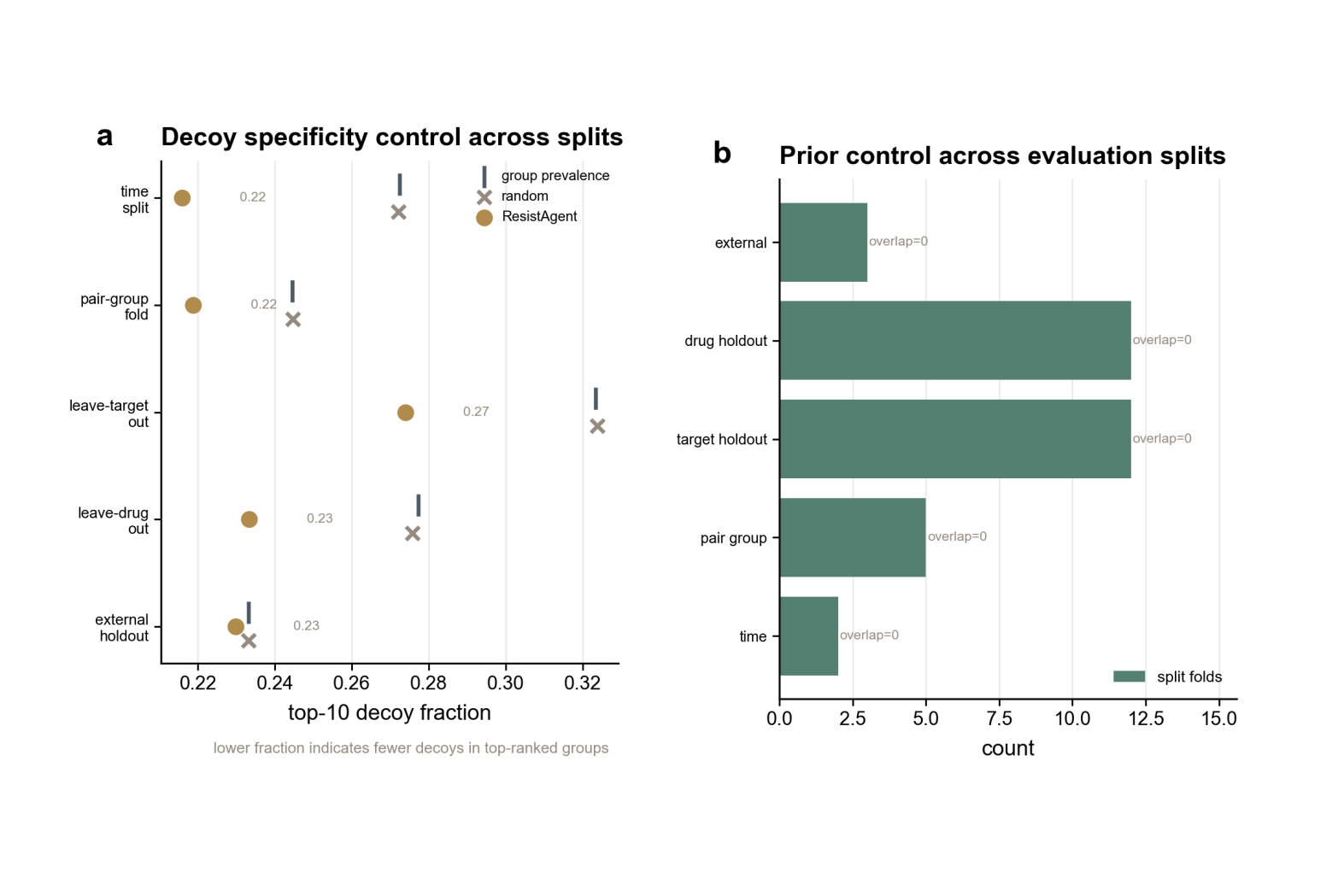

### Supplementary Tables

**Supplementary Table 1. Source databases, evidence tiers and guardrails.**

| **source** | **role** | **table** | **guardrail** |
| --- | --- | --- | --- |
| COSMIC | cancer background/tissue context | master_mutation_table | not used for viral HIV priors |
| DepMap | cancer background context | master_mutation_table | not used for viral HIV priors |
| MdrDB | drug-resistance benchmark rows and observed mutation examples | master_mutation_table | blind benchmark prior excludes MdrDB same-drug signal |
| HIVDB/GenoRx | viral background and drug-selected priors | hiv_*_prior | domain-specific viral prior |

**Supplementary Table 2. Full case overview and template summary.**

| **case_id** | **case_label** | **role** | **site_liability_rows** | **combo_liability_rows** |
| --- | --- | --- | --- | --- |
| egfr_erlotinib | EGFR-Erlotinib | primary | 27 | 3 |
| hiv_rt_rilpivirine | HIV-RT-Rilpivirine | primary | 37 | 16 |
| abl1_nilotinib | ABL1-Nilotinib | boundary | 34 | 0 |

**Supplementary Table 6. Representative robust SAR summary source table.**

| **System** | **Main evaluation unit** | **Representative liabilities captured** | **Dominant structural failure mode(s)** | **Robust design response** | **Quantitative shift** | **Evidence status / caveat** |
| --- | --- | --- | --- | --- | --- | --- |
| EGFR-Erlotinib | site liabilities plus selected observed combos | E746_A750delELREA; T790M; T790M+L858R | pocket rearrangement; anchor/electrostatic shift | retain WT anchors while adding non-hotspot compensation | panel 0.882 vs 0.806; Dep 0.008 vs 0.010 | primary case; short physical support is limited supporting evidence |
| HIV-RT-Rilpivirine | site liabilities plus observed combo panel | E138K; Y181C; V106M+V179D; K101E+Y181C | anchor loss; steric/electrostatic shift; pocket rearrangement | combo-aware buffering and redistributed contact dependence | panel 0.722 vs 0.841; Dep 0.126 vs 0.187 | primary case; robust changes dependence profile with case-specific panel behavior |
| ABL1-Nilotinib | site-only, uncertainty-heavy boundary case | T315I; Y253H; E255K | weaker calibration; uncertainty-heavy structural scoring | boundary characterization for validation-boundary interpretation | panel 0.314 vs 0.297; Pearson r 0.345 | boundary context for interpreting case-dependent generality |

Supplementary Tables 3-5 and 7-8 are provided as separate editable sheets in ResistAgent_Supplementary_Data.xlsx because the full source tables are too large for readable Word layout.

**Supplementary Table 9. Ranking benchmark configuration used to define the validation envelope.**

| **Setting** | **Value** |
| --- | --- |
| model | xgb_dual |
| classifier_weight | 0.75 |
| regression_z_weight | 0.15 |
| docking_z_weight | 0.10 |
| prior_delta_weight | 0.00 |
| time_split | ready |
| folds | 2 |
| interpretation | ranking-first benchmark support within the stated validation envelope |

**Supplementary Table 10. Complete ABL1-Nilotinib boundary-case result summary.**

| **Metric** | **Value** |
| --- | --- |
| case role | boundary / uncertainty-heavy case |
| site liabilities | 34 |
| combo liabilities | 0 |
| dock-vs-MM/GBSA Pearson r | 0.345 |
| high-uncertainty mechanism calls | 21 |
| candidate count | 174 |
| overall panel passing rate | 0.306 |
| robust panel passing rate | 0.314 |
| naive panel passing rate | 0.297 |
| robust top-20 RobustScore median | 0.365 |
| naive top-20 RobustScore median | 0.345 |
| robust top-20 dependence median | 0.152 |
| naive top-20 dependence median | 0.160 |
| evidence-confidence score | 0.568 |
| interpretation | boundary context for case-dependent generality |

**Supplementary Table 11. Parameter and reproducibility configuration used to interpret equations and paired searches.**

| **Setting** | **Value** |
| --- | --- |
| paired primary-case seeds | 101, 202, 303 for EGFR and HIV |
| paired search rounds | 6 |
| proposal_count | 16 |
| beam_width | 6 |
| max_parallel_candidates | 6 |
| target_parallel_workers | 8 |
| search_rank_jitter | 0.35 |
| CVaR lower-tail q | 0.20 |
| robust objective weights | WT 0.8; site 1.0; combo 1.2; alternative anchors 0.5; non-hotspot support 0.35 |
| penalty weights | hotspot fraction 0.25; evidence uncertainty 0.25; synthesis penalty 0.25 |
| evidence-confidence weights | calibration 0.35; sample support 0.15; target coverage 0.15; target count 0.15; uncertainty/missingness terms 0.20 |
| panel selection | cumulative risk-mass selection with predefined minimum and maximum panel sizes; HIV retains a dedicated observed-combo budget |
| chemistry constraints | chemistry engine v2; synthesis realism on; dynamics_lite off; dep_focus on; scaffold_tail_diversity on |
| large-source tables | full mutation ranks, mutation effects, top designed molecules and full benchmark tables are supplied as editable workbook sheets |
